# Modelling microbiome recovery after antibiotics using a stability landscape framework

**DOI:** 10.1101/222398

**Authors:** Liam P. Shaw, Hassan Bassam, Chris P. Barnes, A. Sarah Walker, Nigel Klein, Francois Balloux

## Abstract

Treatment with antibiotics is one of the most extreme perturbations to the human microbiome. Even standard courses of antibiotics dramatically reduce the microbiome’s diversity and can cause transitions to dysbiotic states. Conceptually, this is often described as a ‘stability landscape’: the microbiome sits in a landscape with multiple stable equilibria, and sufficiently strong perturbations can shift the microbiome from its normal equilibrium to another state. However, this picture is only qualitative and has not been incorporated in previous mathematical models of the effects of antibiotics. Here, we outline a simple quantitative model based on the stability landscape concept and demonstrate its success on real data. Our analytical impulse-response model has minimal assumptions with three parameters. We fit this model in a Bayesian framework to data from a previous study of the year-long effects of short courses of four common antibiotics on the gut and oral microbiomes, allowing us to compare parameters between antibiotics and microbiomes, and further validate our model using data from another study looking at the impact of a combination of last-resort antibiotics on the gut microbiome. Using Bayesian model selection we find support for a long-term transition to an alternative microbiome state after courses of certain antibiotics in both the gut and oral microbiomes. Quantitative stability landscape frameworks are an exciting avenue for future microbiome modelling.

## Introduction

### Stability and perturbation in the microbiome

The human microbiome is a complex ecosystem. While stability is the norm in the gut microbiome, disturbances and their consequences are important when considering its impact on health (1). A course of antibiotics is a major perturbation, typically leading to a marked reduction in species diversity before subsequent recovery (2). Aside from concerns about the development of antibiotic resistance, even a brief course can result in long-term effects on community composition (3). However, modelling the recovery of the microbiome is challenging, due to the difficulty of quantifying the *in vivo* effects of antibiotics on the hundreds of co-occurring species that make up microbial communities within the human body.

Artificial perturbation experiments are widely used to explore the underlying dynamics of macro-ecological systems (4). In the context of the gut microbiome, the effects of antibiotics have previously been investigated (5–7). However, despite interest in the application of ecological theory to the gut microbiome (8), the nature of this recovery after antibiotics remains unclear. While responses can appear highly individualized (7) this does not preclude the possibility of generalized models applicable at the population level.

Applying mathematical models to other ecological systems subject to perturbation can give useful insight (9–11). Crucially, it allows the comparison of different hypotheses about the system using model selection. Developing a consistent mathematical framework for quantifying the long-term effects of antibiotic use would facilitate comparisons between different antibiotics and different regimens, with the potential to inform approaches to antibiotic stewardship (12).

### Previous modelling approaches

A great deal of modelling work has focused on the gut microbiome’s response to antibiotic perturbation. We mention a few important examples here. Bucci et al. (13) used a two-compartment density model with species categorised as either antibiotic-tolerant or antibiotic-sensitive, and fitted their model to data from Dethlefsen and Relman (7). In a later review, Bucci and Xavier argued that models of wastewater treatment bioreactors could be adapted for the gut microbiome, with a focus on individual-based models (14). The most commonly used individual-based model is the multispecies Generalized Lotka-Volterra (GLV) model, which describes pairwise interactions between bacterial species (or other groupings). In a pioneering work, Stein et al. (15) extended a GLV model to include external perturbations, and fitted their model to a study where mice received clindamycin and developed *Clostridium difficile* infection (CDI) (16). The same approach was also subsequently applied to human subjects in, identifying a probiotic candidate for treating CDI (17). Bucci et al. (18) have combined and extended their previous work into an integrated suite of algorithms (MDSINE) to infer dynamical systems models from time-series microbiome data.

While all of these models have provided useful insights into microbiome dynamics, to make meaningful inference they require dense temporal sampling and restriction to a small number of species. For example, the examples of application of MDSINE had “26–56 time points” for accurate inference of dynamics, measurements of relative concentrations of bacteria, and frequent shifts of treatment — for these reasons the *in vivo* experiments were conducted in gnotobiotic mice (18). Similarly, Stein et al. restricted their analysis of CDI to the ten most abundant bacterial genera (15). Such restrictions reduce the suitability of these methods for opportunistic analysis of existing 16S rRNA gene datasets from the human microbiome, which currently comprise the majority of clinically relevant datasets. GLV models can undoubtedly be extremely useful for simple synthetic consortia, as shown by Venturelli et al. who inferred the dynamics of a 12-species community (19). However, it has been shown that even for very small numbers of species, pairwise microbial interaction models do not always accurately predict future dynamics, suggesting that pairwise modelling has its own limitations (20).

Starting from broader ecological principles allows quantitative investigation of high-level statements and hypotheses about microbiome dynamics. For example, Coyte et al. built network models based on principles from community ecology to show that competitive interactions in the gut microbiome are associated with stable states of high diversity (21). More recently, Goyal et al. took inspiration from the ‘stable marriage problem’ in economics and showed that multiple stable states in microbial communities can be explained by nutrient preferences and competitive abilities (22). There is therefore great value in exploring alternative modelling approaches to GLV models as well as continuing to refine and extend them.

### A stability landscape approach

In a popular schematic picture taken from classical ecology, the state of the gut microbiome is represented by a ball sitting in a stability landscape (1,23–25). Perturbations can be thought of as forces acting on the ball to displace it from its equilibrium position (25) or as alterations of the stability landscape (26). While this image is usually provided only as a conceptual model to aid thinking about the complexity of the ecosystem, we use it here to derive a mathematical model.

We model the effect of a brief course of antibiotics on the microbial community’s phylogenetic diversity as the impulse response of an overdamped harmonic oscillator (Figure 1; see Methods), and compare parameters for four widely-used antibiotics by fitting to empirical data previously published by Zaura et al. (3). This model is significantly less complicated than previous models developed for similar purposes, but still captures some of the essential emergent features of such a system while avoiding the computational difficulties of fitting hundreds of parameters to a sparse dataset. After demonstrating the effectiveness of this modelling approach for the gut and oral microbiome, we also show that the framework can easily be used to test hypotheses about microbiome states. We compare a model variant which allows a transition to a new equilibrium, and find that this model is better supported for clindamycin and ciprofloxacin, allowing us to conclude that these antibiotics can produce state transitions across different microbiomes. We also find a transition to a new equilibrium when reanalyzing a separate dataset from Palleja et al. (27), using a different diversity metric that had already been calculated by the authors (richness) based on applying a different sequencing approach (shotgun metagenomics) to an different antibiotic regimen (meropenem, gentamicin and vancomycin). This modelling approach can therefore be easily applied to sparse datasets from different human microbiomes, antibiotics, and sequencing approaches, providing a simple but consistent foundational framework for quantifying the *in vivo* impacts of antibiotics.

**Figure 1.**
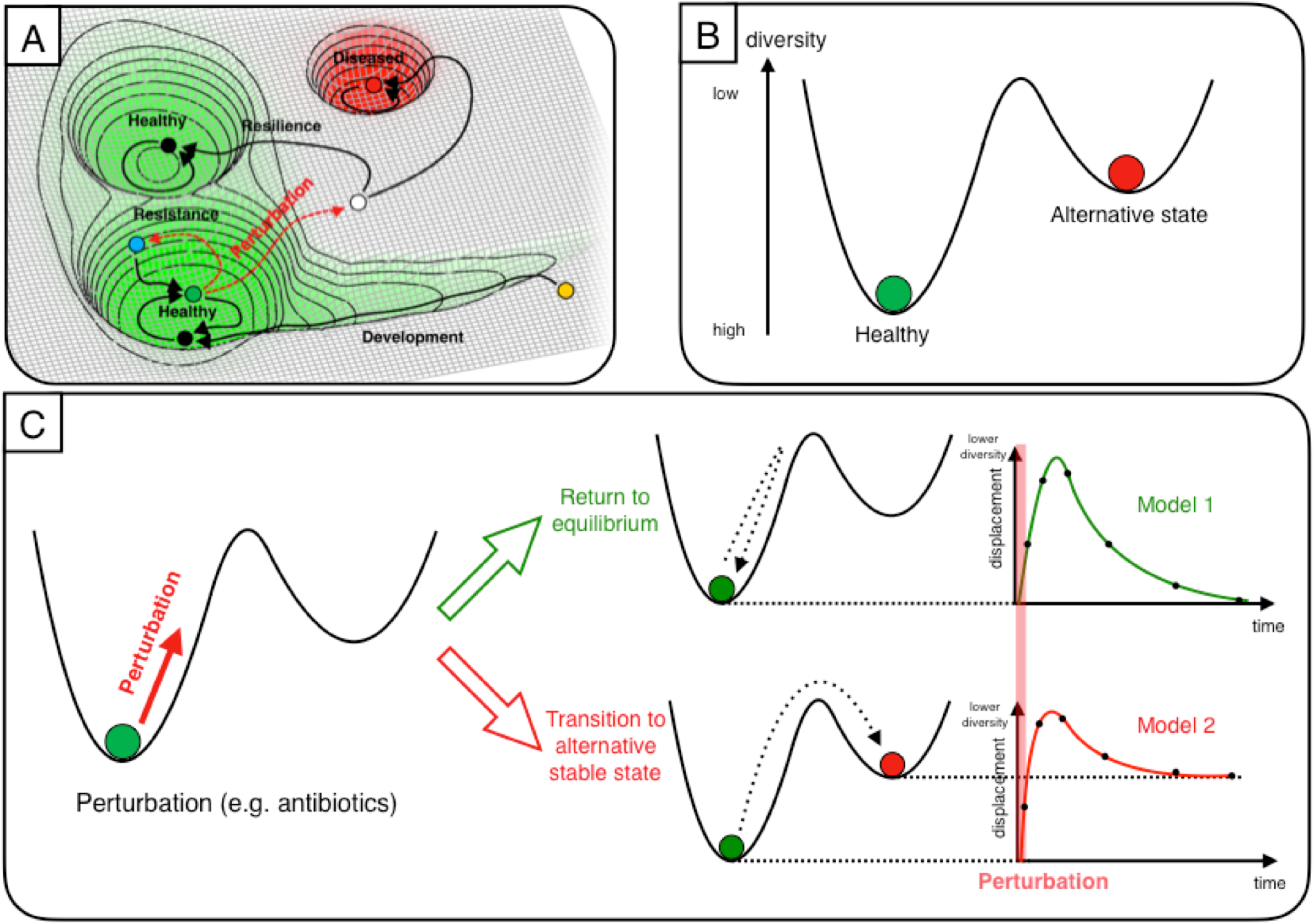
A stability landscape framework for antibiotic perturbation to the microbiome. We represent the gut microbiome as a unit mass on a stability landscape, where height corresponds to phylogenetic diversity. (A) The healthy human microbiome can be conceptualised as resting in the equilibrium of a stability landscape of all possible states of the microbiome. Perturbations can displace it from this equilibrium value into alternative states (adapted from Lloyd-Price et al. (25)). (B) Choosing to parameterise this stability landscape using diversity, we assume that there are just two states: the healthy baseline state and an alternative stable state. (C) Perturbation to the microbiome (e.g. by antibiotics) is then modelled as an impulse, which assumes the duration of the perturbation is short relative to the overall timescale of the experiment. We consider the form of the diversity time-response under two scenarios: a return to the baseline diversity; and a transition to a different value of a diversity (i.e. an alternative stable state).

## Results

### Ecological theory motivates a simplified representation of the microbiome

Taking inspiration from classical ecological theory, the microbiome can be considered as an ecosystem existing in a stability landscape: it typically rests at some equilibrium, but can be displaced (Figure 1A). The shape of this landscape is determined by the extrinsic environmental factors – a change in these could produce a (reversible) change in the shape of the landscape e.g. shifts in the human gut microbiome when an individual temporarily relocates from the USA to Southeast Asia, which are reversed on return (28). Here we assume that these environmental factors remain approximately constant at the timescales considered (< 1 year), based on previous observations e.g. that the oral microbiome can exhibit stability over periods of at least three months (29), and the gut microbiome for up to five years (30). Any quantitative model of the microbiome based on this concept then requires a definition of equilibrium and displacement. While earlier studies sought to identify a equilibrium core set of ‘healthy’ microbes, disturbances of which would quantify displacement, it has become apparent that this is not a practical definition due to high inter-individual variability in taxonomic composition (25). More recent concepts of a healthy ‘functional core’ appear more promising, but characterization is challenging, particularly as many gut microbiome studies use 16S rRNA gene sequencing rather than shotgun sequencing.

For these reasons, we choose a metric that offers a proxy for the general functional potential of the gut microbiome: phylogenetic diversity (25). Diversity is commonly used as a summary statistic in microbiome analyses and higher diversity in the gut microbiome has previously been generally associated with health (31) and temporal stability (32). Of course, describing the microbiome using only a single number loses a great deal of information. Furthermore, there is no general relationship between ‘diversity’ and ‘stability’: the relationship depends on the specific definitions of both quantities. We are interested in the scenario of fixed-length perturbations, where ‘stability’ can be quantified in terms of the rate of return to equilibrium after perturbation – this stability has been empirically observed to increase with greater species diversity (33). If we are seeking to build a general model of microbiome recovery after perturbation, it seems appropriate to consider a simple metric first to see how such a model performs before developing more complicated definitions of equilibrium, which may generalise poorly across different niches and individuals. Other scenarios might require different definitions and correspondingly different assumptions.

We assume the equilibrium position to have higher diversity than the points immediately surrounding it i.e. creating a potential well (Figure 1B). However, there may be alternative stable states (Figure 1B) which perturbations may move the microbiome into (Figure 1C). These states may be either higher or lower in diversity; for our purposes, all we assume is that they are separated from the initial equilibrium by a potential barrier of diversity i.e. a decrease of diversity is required to access them, which helps to keep the microbiome at equilibrium under normal conditions. Our strong assumption here is obviously influenced by the specific perturbation scenario: we know *a priori* that antibiotics decrease diversity in the short-term, so we are not attempting to model other types of transition. However, considering the empirical evidence and the consensus view among ecologists that a diversity of species with a range of sensitivities to different environmental conditions should lead to greater stability (33,34), we feel this rule may hold in a range of other scenarios: ecosystems may often exist at or near a (local) maximum value of diversity. An equilibrium does not have to be believed in as a true and eternal state; we attempt here to use insights from theoretical communities with well-defined mathematical equilibria to move to modelling empirical communities where they are approximate concepts, valid over some given time period.

### The model

Mathematically, small displacements of a mass from an equilibrium point can be approximated as a simple harmonic oscillator (35) for any potential function (continuous and differentiable). This approximation comes naturally from the first terms in the Taylor expansion of a function (36), and can be extremely accurate for small perturbations. By assuming the local stability landscape of the microbiome can be reasonably approximated as a harmonic potential, we are assuming a ‘restoring’ force proportional to the displacement x from the equilibrium position (-*kx*) and also a ‘frictional’ force acting against the direction of motion ddd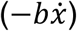. The system is a damped harmonic oscillator with the following equation of motion:

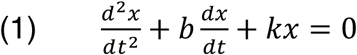

Additional forces acting on the system — perturbations — appear on the right-hand side of this equation. Consider a course of antibiotics of duration *τ*. If we are interested in timescales of *T* ≫ *τ* (e.g. the long-term recovery of the microbiome a year after a week-long course of antibiotics) we can assume that this perturbation is of negligible duration. This assumption allows us to model it as an impulse of magnitude *D* acting at time *t* = 0:

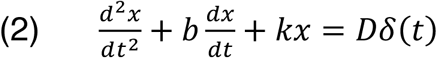

This second-order differential equation can be solved analytically and reparameterised (see Methods) to give a model with three parameters, with a general qualitative shape of sudden change followed by slower recovery (Model 1, Figure 1C):

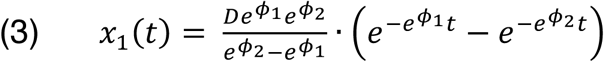

### Fitting the model to empirical data

We fit the model to published data from a paper from Zaura et al. (3) where individuals received a ten-day course of either a placebo or one of four commonly-used antibiotics (Table 1). Faecal and saliva samples were taken at baseline (i.e. before treatment), then subsequently directly after treatment, then one month, two months, four months, and one year after treatment. Zaura et al. conducted pairwise comparisons between timepoints and comprehensively reported statistical associations, but did not attempt any explicit modelling of the time-response over the year. This dataset provides an ideal test case for our model. Not only does it allow us to simultaneously model the recovery of both the gut and oral microbiomes after different antibiotics, but it also demonstrates how our modelling framework permits further conclusions beyond the scope of the initial study. We additionally fit our model to a shotgun metagenomics dataset from a recent antibiotic perturbation experiment from Palleja et al. (27).

**Table 1.** Number of individuals in each treatment group. Only individuals with a complete set of 6 samples with >1,000 reads in each were retained for model fitting. For demographic characteristics of the complete treatment groups see Table 1 of Zaura et al. (3).

| Antibiotic treatment group | *n* (gut microbiome) | *n* (oral microbiome) |
| --- | --- | --- |
| Placebo | 22 | 21 |
| Ciprofloxacin | 9 | 9 |
| Clindamycin | 9 | 9 |
| Minocycline | 10 | 10 |
| Amoxicillin | 12 | 12 |

### A stability landscape framework successfully describes initial microbiome dynamics

Although individual microbiome responses were personalized (Supplementary Figure 1), normalized diversity trajectories followed the same general pattern for each antibiotic (Supplementary Figure 2). We used a Bayesian approach to fit the model to each treatment group and microbiome separately, successfully capturing the main features of the initial response to antibiotics (Figure 2). Diversity decreased (i.e. displacement from equilibrium increased) before a slow return to equilibrium. Despite large variability between samples from the same treatment group, reassuringly the placebo group clearly did not warrant an impulse response model, whereas data from individuals receiving antibiotics was qualitatively in agreement with the model. Even without the model, it is apparent that clindamycin and ciprofloxacin represent greater disturbances to the microbiome than minocycline and amoxicillin, but a consistent model allows comparison of the values of various parameters (see below).

**Figure 2.**
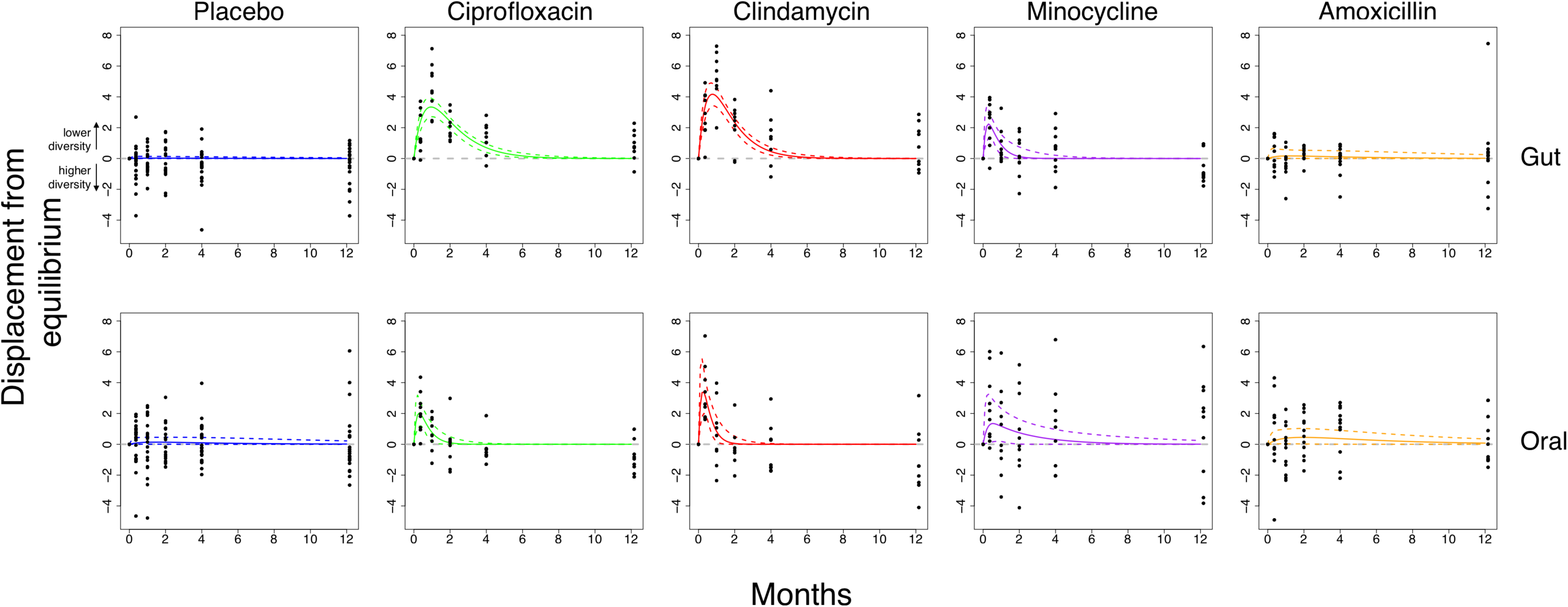
The model captures the dynamics of recovery for the gut and oral microbiomes after antibiotics. Bayesian fits for participants taking either a placebo (blue; n=21/22 for gut/oral), ciprofloxacin (green; n=9), clindamycin (red; n=9), minocycline (purple; n=10), and amoxicillin (orange; n=12). The mean phylogenetic diversity from 100 bootstraps for each sample (black points) and median and 95% credible interval from the posterior distribution (bold and dashed coloured lines, respectively). The grey line indicates the equilibrium diversity value, defined on a per-individual basis relative to the mean baseline diversity. The biased skew of residuals after a year in certain treatment groups suggests the possibility of a transition to an alternative stable state with a different value of diversity.

In their original analysis, Zaura et al. noted significantly (*p* < 0.05) reduced Shannon diversity in individuals receiving ciprofloxacin comparing samples after a year to baseline. This reduced diversity could in principle merely be due to slow reconstitution and return to the original equilibrium. However, by taking into account each individual’s temporal response with a model rather than pairwise comparisons it appears that slow reconstitution cannot be the whole story. Instead, the skewed distribution of residuals after a year, when the response has flattened off, indicates that the longer-term dynamics of the system do not obey the same impulse response as the short-term dynamics. A scenario involving a long-term transition to an alternative stable state is consistent with this observation (Figure 1). We therefore developed a variant of the model to take into account alternative equilibria, aiming to test the hypothesis that the microbiome had transitioned to an alternative stable state.

### Support for the existence of antibiotic-induced state transitions

In our approach, a transition to an alternative stable state means that the value of diversity displacement from the original equilibrium asymptotically tends to a non-zero value. There are many options for representing this mathematically; for reasons of simplicity, we add a single parameter *A* and a term that asymptotically grows over time (Model 2, Figure 1C):

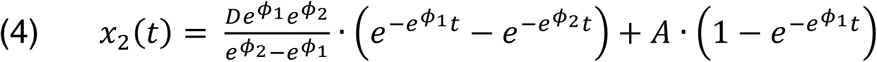

Qualitatively, this slightly more complex model gave a similar fit (Figure 3) but some treatment groups had a clear non-zero final displacement from equilibrium, corresponding to an alternative stable state. We compared models with the Bayes factor *BF*, where *BF* > 1 indicates support for a state transition (Table 2). A state transition was supported in the ciprofloxacin and clindamycin treatment groups for both the gut (*BF*_*cipro*_ = 3.06, *BF*_*clinda*_ = 10.94) and oral (*BF*_*cipro*_ = 16.87, *BF*_*clinda*_ = 7.47) microbiomes. The posterior estimates for the asymptote parameter in the gut microbiome were positively skewed (Figure 4), providing evidence of a transition to a state with lower phylogenetic diversity. Contrastingly, in the oral microbiome the asymptote parameter was negatively skewed, suggesting a transition to a state with greater phylogenetic diversity. Strikingly, these are the respective states associated with poorer health in both microbiomes.

**Table 2.** Bayes factors for model comparisons for each antibiotic group. The Bayes factor (BF) allows model selection, here for model 2 (with a state transition) against model 1 (no state transition). Following Kass and Raffery, we interpret BF>3 as positive evidence in favour of model 2 (51).

| Antibiotic treatment group | Gut microbiome | Oral microbiome |
| --- | --- | --- |
| Ciprofloxacin | **3.06** | **16.87** |
| Clindamycin | **10.94** | **7.47** |
| Minocycline | 2.11 | 2.42 |
| Amoxicillin | 1.51 | 1.31 |

**Figure 3.**
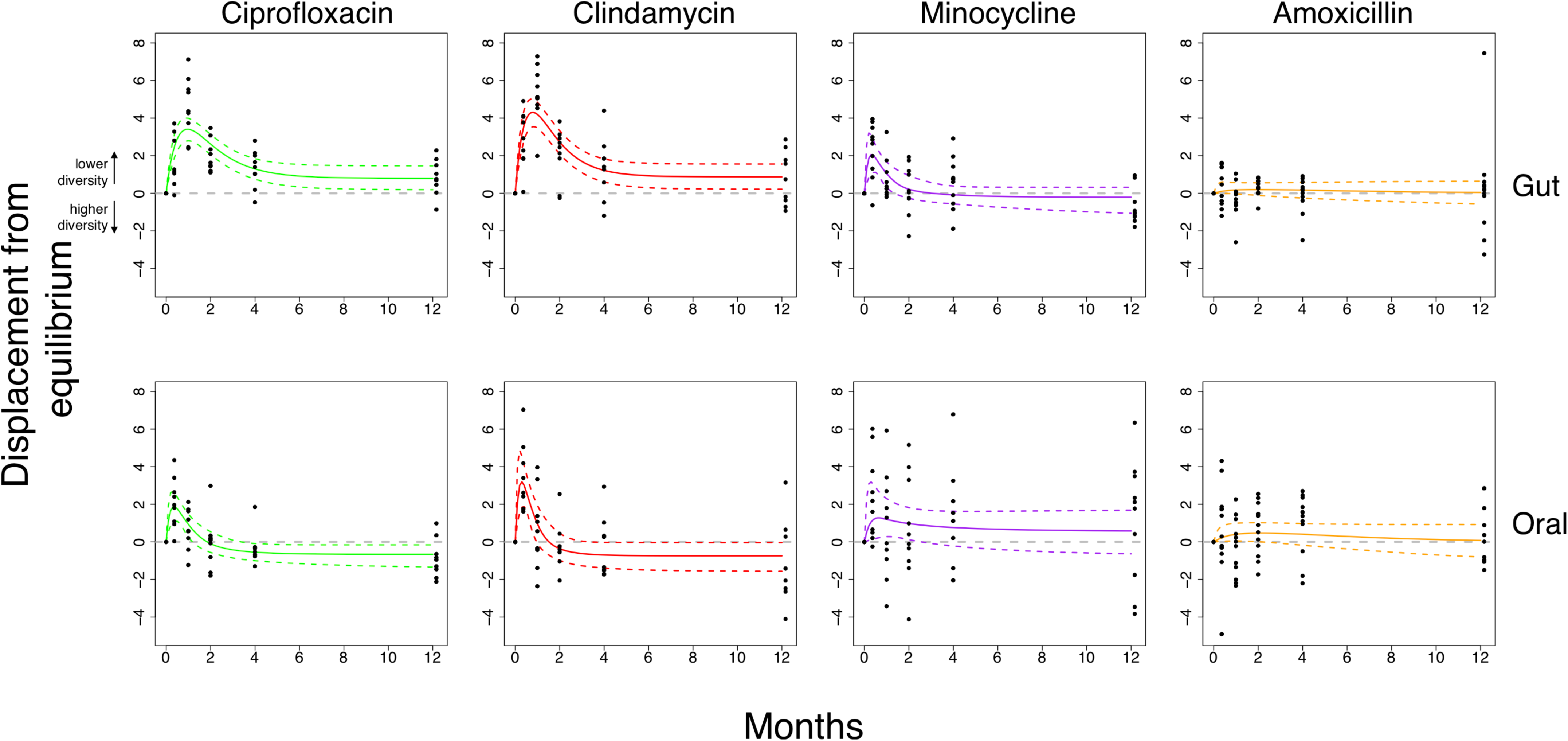
A model with a possible state transition is better supported for clindamycin and ciprofloxacin. Bayesian fits for participants taking either ciprofloxacin (green; n=9), clindamycin (red; n=9), minocycline (purple; n=10), and amoxicillin (orange; n=12). The mean phylogenetic diversity from 100 bootstraps for each sample (black points) and median and 95% credible interval from the posterior distribution (bold and dashed coloured lines, respectively). The grey line indicates the equilibrium diversity value, defined on a per-individual basis relative to the mean baseline diversity. The non-zero-centred asymptotes indicates support for a state transition in both the gut and oral microbiomes after ciprofloxacin and clindamycin. See Table 2 for Bayes Factors comparing model 2 to model 1.

**Figure 4.**
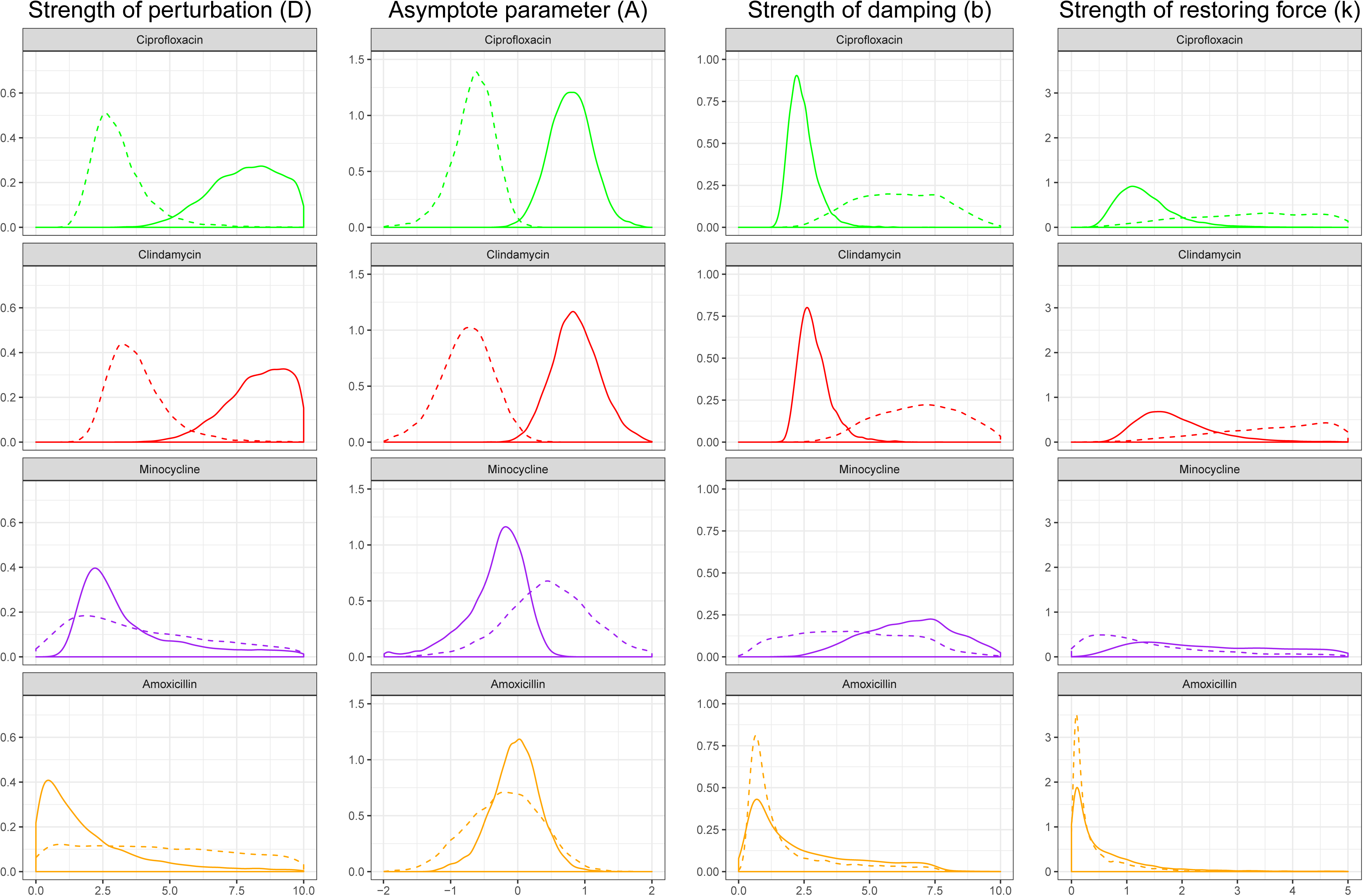
Posterior parameter estimates for model with a possible transition to an alternative stable state. The posterior distributions from Bayesian fits of model 2 (eq. 7) to empirical data from the gut (solid) and oral microbiomes (dashed) of individuals who received ciprofloxacin (green), clindamycin (red), minocycline (purple), and amoxicillin (orange). The posterior probability distribution is a way of visualizing the uncertainty in parameter values after model fitting (a tighter peak indicates more certainty about the parameter value), and can be subsequently used to derive e.g. interval estimates. Because the sum under the distribution is defined as being equal to one, the scale of the y-axis depends on the range of the x-axis i.e. it has no absolute meaning.

### Comparison of parameters between antibiotics

Comparing the posterior distribution of parameters for model 2 fits allows quantification of ecological impact of different antibiotics (Table 3, Figure 4). Unsurprisingly, greater perturbation is correlated with the transition to an alternative stable state. We can also consider the ecological implications of the parameters we observe. The damping ratio 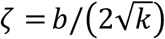 summarises how perturbations decay over time, and is considered as an inherent property within the model. Therefore, if our modelling framework and ecological assumptions were valid we would expect to find a consistent damping ratio across both the clindamycin and ciprofloxacin groups in the gut microbiome. This is indeed what we observed with median (95% credible interval) damping ratios of *ζ*_clinda_=1.07 (1.00-1.65) and *ζ*_cipro_=1.07 (1.00-1.66), substantially different from the prior distribution, supporting the view that the gut microbiome dynamics can be approximated by a damped harmonic oscillator.

**Table 3.**
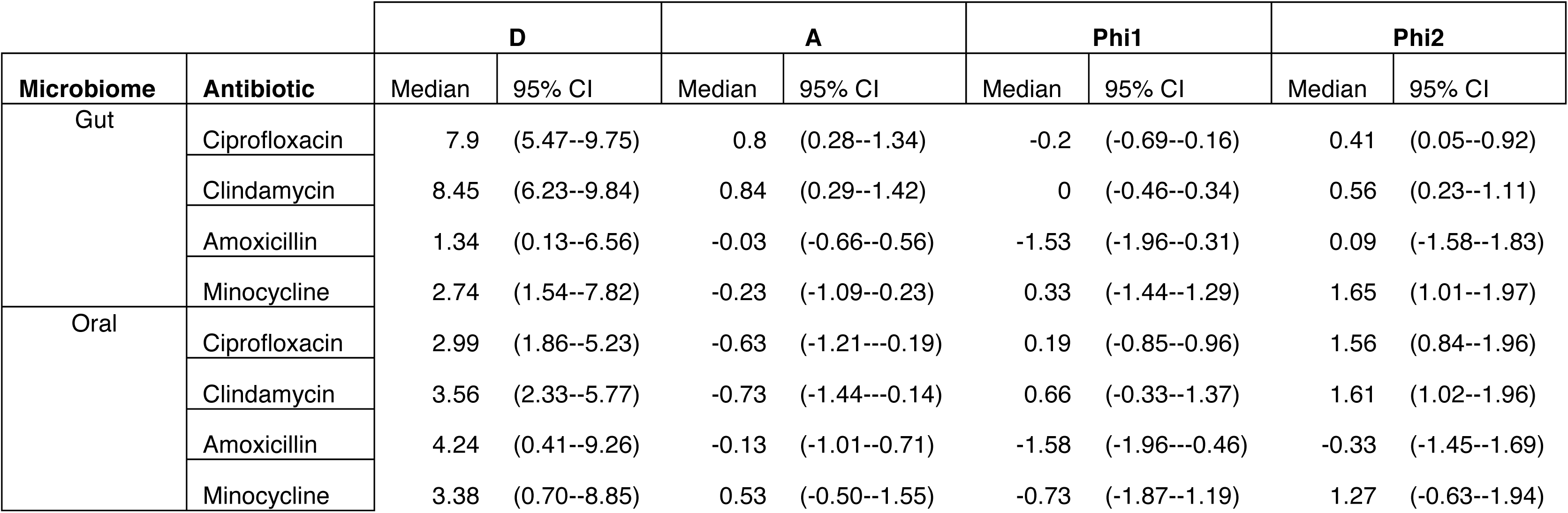
Median and 95% credible intervals for all model parameters for each treatment group. Results from Bayesian fitting of the full model (model 2) to each of the eight possible treatment groups (4 antibiotics x 2 microbiomes).

### Application to other sequencing approaches and diversity metrics

In a recent paper, Palleja et al. (27) used deep shotgun metagenomic sequencing to characterize the gut microbiomes of twelve healthy men for 180 days after a four-day course of meropenem, gentamicin and vancomycin. They observed that while there were no significant differences in the Shannon diversity between baseline and day 180, there was a significant difference in species richness. Furthermore, they noted that compositional changes between day 42 and day 180 were not significantly different when compared to paired samples in Human Microbiome Project controls. Both these two pieces of information suggest a state transition, although Palleja et al. do not use this terminology and instead refer to a return to “stable composition”. Reanalyzing this species richness data with our model (Supplementary File 10) shows that the gut microbiomes of these individuals had transitioned to a new alternative state (Supplementary Figure 3; Table 4), strengthening Palleja et al.’s conclusion that some bacteria may have been permanently lost (27). Other future metagenomic datasets could also be analyzed using our model in this way.

**Table 4.** Median and 95% credible intervals for all model parameters for the Palleja et al. (2018) dataset. Results from Bayesian fitting of the full model (model 2) to the rescaled richness dataset. These should not be directly compared to the fitting to the Zaura et al. dataset, as the diversity metrics used are different. The estimated Bayes factor supporting model 2 over model 1 was *BF* = 75.6.

| D |  | A |  | Phi1 |  | Phi2 |  |
| --- | --- | --- | --- | --- | --- | --- | --- |
| Median | 95% CI | Median | 95% CI | Median | 95% CI | Median | 95% CI |
| 2.24 | (1.02--3.88) | 0.60 | (0.26--0.91) | 0.39 | (-0.37--1.23) | 1.70 | (0.99--1.99) |

### Connection to generalized Lotka-Volterra models

We sought to establish a link between our framework and the conventional ‘bottom-up’ approach of GLV models. We investigated the behaviour of a 3-species Lotka-Volterra system to establish if perturbation to an alternative state was possible in this simple case (see Methods). We found that only 0.079% of 3-species Lotka-Volterra systems exhibit the behaviour required by our two-state model – because we assume that diversity is continuous, this model is unrealistic for small species numbers. However, for larger numbers, theoretical ecology gives strong justification for our assumptions. As the number of species *n* increases, the number of fixed points which are stable increases (37), and the proportion of simulations from random parameters that have multiple fixed points also increases e.g. with *n* = 400, this proportion is >97% (38). This suggests that the overwhelming majority of *mathematically possible* systems at relevant numbers of species exhibit multiple fixed points; the fraction of *biologically possible* systems exhibiting this behaviour is likely even higher. Furthermore, when the realistic assumption of resource competition is incorporated, all these fixed points become stable or marginally stable (38). Goyal et al. recently showed that multiple resilient stable states can exist in microbial communities if microbes utilize nutrients one at a time (22). We can therefore state confidently that: the gut microbiome can be treated as having multiple stable equilibria; its community composition is history-dependent; and perturbations lead to transitions between the multiple possible stable states. These assumptions are the basis of the simplistic coarse-grained model we describe here, which effectively takes these high-level emergent properties of multi-species Lotka-Volterra models to build a substantially simpler model based on a single, commonly-used metric: diversity.

## Discussion

Starting from a conceptual picture of the microbiome resting in a stability landscape, we have developed a mathematical model of its response to antibiotic perturbation. Our framework, based on phylogenetic diversity, successfully captures the dynamics of a previously published 16S rRNA gene dataset for four common antibiotics (3), providing quantitative support for these simplifying ecological assumptions. Using model selection, our framework provides additional insight — we find that the effects of clindamycin persist for a year after exposure, and also identify a state transition in the oral microbiome with clindamycin, neither of which was not detected by the initial authors. We also demonstrate the flexibility of our model by fitting it to data from a recent shotgun metagenomics antibiotic perturbation experiment (27).

While pairwise comparisons of diversity can still identify differences in microbiome state, they provide no information on dynamics. Our framework therefore gives additional insight in this regard. Zaura et al. observed that the lowest diversity in the gut microbiome was observed after a month rather than immediately after treatment (3). This cannot be due to a persistence of the antibiotic effect, as all antibiotics used have short half-lives of the order of hours (39,40). Within our framework, this can be understood as an overdamping effect: we found a consistent damping ratio for both ciprofloxacin and clindamycin. This is not in itself a complete explanation (it does not identify a mechanistic biological process behind the ‘damping’) but it gives us insight that such a process may exist.

We have also demonstrated how our framework could be used to compare different hypotheses about the long-term effects of antibiotic perturbation by fitting different models and using Bayesian model selection. Our model provides an additional line of evidence that while short-term restoration obeys a simple impulse response, the underlying long-term community state can be fundamentally altered by a brief course of antibiotics, as suggested previously (7), raising concerns about the long-term impacts of antibiotic use. While this state transition may not necessarily equate to any negative health impacts for the host (none of the participants involved in the original study (3) reported any gastrointestinal disturbance), the transition to a new state with reduced diversity in the gut microbiome may increase the risk of colonisation and overgrowth of pathogenic species. It has been shown experimentally in *Drosophila* that invasion and subsequent colonization is a stochastic process that is reduced by a more diverse gut (41). Conversely, in the oral microbiome the state transition was to a state with *increased* diversity, which is correspondingly associated with a greater risk of disease in the oral cavity (42). We believe this makes sense within a stability landscape framework. Even if only marginal, when considered at a population level, such effects may mean that antibiotics have substantial negative health consequences, which could support reductions in the length of antibiotic courses independently of concerns about antibiotic resistance (43). Modelling the long-term impact on the microbiome of different doses and courses could help to influence antibiotic use in routine clinical care. Our sample size is small, so the precise posterior estimates for parameters that we obtain should not be over-interpreted, but comparing antibiotics using these parameter estimates represents another practical application.

Our framework lends itself naturally to comparing different dynamical models. We see our two variant models as a starting point for stability landscape approaches, and would hope that better models can be constructed. Hierarchical mixed effects models may offer an improved fit, particularly if they take into account other covariates; we lacked the necessary metadata on the participants from the original study (Table 1) to explore such models. Furthermore, diversity as a single metric clearly fails to capture all the complexity of the microbial community and its interactions, and there are multiple issues with calculating it accurately. Nevertheless, the observation that treating phylogenetic diversity as the key variable in the stability landscape captures microbiome dynamics supports observations of functional redundancy in the microbiome (31). An interesting extension of this work would be to systematically fit the model to a variety of diversity metrics or other summary statistics and assess the model fit to see which metric (or combination of metrics) is most appropriately interpreted as the state variable parameterising the stability landscape. A complementary approach could consider the ‘resistome’, which should conversely rise in diversity after antibiotic treatment (44).

Expanding the stability landscape approach to other microbiome datasets might require re-examining our assumptions. A non-diversity metric might well be superior in situations where the ‘ideal’ state of the microbiome can be clearly defined: for example, in a bioreactor where experimental conditions can be maintained to consistently produce a given equilibrium community composition, one could define displacement using ecological distance from this well-defined state. However, even a single microbial strain under constant conditions can exhibit astonishing and unpredictable complexity over time (45), so we feel that diversity is a good starting point. As a general guide for experiments, because an impulse response model for a perturbation of length *τ* assumes an experiment length *T* ≫ *τ*, we suggest the appropriate sampling effort for microbiomes undergoing perturbation should be to have samples immediately before and after, as well as samples out to *T* > 30*τ*. We recommend more dense sampling at longer times to firmly establish whether a new equilibrium state has been reached.

We would not expect the behaviour of the microbiome after longer or repeated courses of antibiotics to be well-described by our model, which assumes a course of negligible duration. Nevertheless, it would be possible to use the stability landscape framework given here to obtain an analytic form for the possible system response by convolving any given perturbation function with the impulse response, although the model might break down in such circumstances. If the model fails to give a good fit, the inclusion of higher-order terms in the Taylor expansion of the chosen potential function about equilibrium (and correspondingly modifying eq. 1) could prove useful.

As we have demonstrated, while the precise nature of the gut microbiome’s response to antibiotics is individualized, a general model still captures important dynamics. We believe it would be a mistake to assume that our model is ‘too simple’ to provide insight on a complex ecosystem. At this stage of our understanding, creating a comprehensive inter-species model of the hundreds of members of the gut microbiome is intractable; it may also be unnecessary if the aim is to inform clinical treatment based on sparse data. We believe there is a place for both fine-grained models using pairwise interactions — particularly for systems of reduced complexity — and coarse-grained models built from high-level ecological principles, as we have demonstrated here. Shade has argued that microbiome diversity is “the question, not the answer” (46); diversity can still be an extremely useful quantity for modelling, but the patterns we report here warrant more mechanistic investigation. We have argued that a ‘top-down’ framework with multiple stable states of different diversities is consistent with the emergent behaviour of a multispecies Lotka-Volterra model. Further mathematical work to connect these two extremes would be worthwhile.

## Supporting information

Supplementary information - description of files

## Acknowledgements

LPS was supported by the Engineering and Physical Sciences Research Council [EP/F500351/1] and the Reuben Centre for Paediatric Virology and Metagenomics. CPB is supported by the Wellcome Trust [209409/Z/17/Z].We are grateful to Zaura et al. (3) and Palleja et al. (27) for making their data openly available, enabling the reanalysis with our modelling framework presented here.

## Authors’ contributions

LPS conceived the model, performed analyses, and wrote the paper. LPS, CPB, HB, and FB conceived the analysis of the Lotka-Volterra system, which was performed by HB. All authors contributed to the discussion and development of the model, gave comments, and read and approved the final manuscript.

## Competing interests

The authors declare that they have no competing interests.

## Data availability

The original sequencing dataset from Zaura et al. (3) used in this paper is available in the Short Read Archive (SRA Accession: <u>SRP057504</u>). Full code and reanalyzed datasets supporting the conclusions of this article are included as Supplementary Information. A full archive of main analyses including cached model fits is available in figshare (https://figshare.com/s/d62d6e90f96dc63c2769.)

## Materials and methods

### Mathematical model of trajectories in the potential landscape

Treating the microbiome as a unit mass resting in a stability landscape parameterised by phylogenetic diversity leads to a second-order differential equation. To solve this equation, we assume that *b*^2^ > 4*k* (the ‘overdamped’ case) based on the lack of any oscillatory behaviour previously observed in the microbiome. Then, subject to the initial conditions *x*(*t* = 0) = 0 and 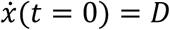 we obtain the following equation describing the system’s trajectory:

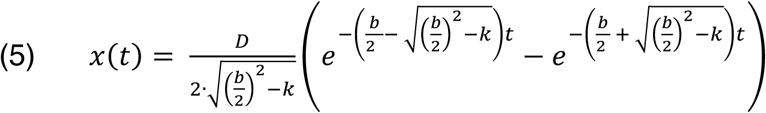

Fitting the model therefore requires fitting three parameters: *b* (the damping on the system), *k* (the strength of the restoring force), and *D* (how strong the perturbation is). For the purposes of fitting the model, where parameters having allowed values of real numbers (-∞, +∞) is preferable to only positive numbers (0, +∞), we choose to reparameterise the model using the following definitions:

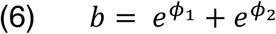

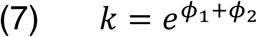

This results in Model 1 (eq. 3, Figure 1C).

Antibiotics may lead not just to displacement from equilibrium, but also state transitions to new equilibria (2). To investigate this possibility, we also consider a model where the value of equilibrium diversity asymptotically tends to a new value *A*. To minimise model complexity, we add a single parameter with a term that asymptotically grows over time, resulting in Model 2 (eq. 4, Figure 1C).

### Main experimental dataset

To apply our framework, we fitted both models to two empirical datasets (see Supplementary Text 1). Zaura et al. (3) conducted a study on the long-term effect of antibiotics on the gut and oral microbiomes, where individuals were randomly assigned to one of five treatment groups: placebo, clindamycin, ciprofloxacin, minocycline, or amoxicillin (Table 1). Samples were collected at baseline (before treatment), immediately after exposure, then at one month, two months, four months, and one year after treatment.

### Phylogenetic diversity

There are many possible diversity metrics that could be used to compute the displacement from equilibrium. Because of our assumptions (see ‘Ecological theory motivates a simplified representation of the microbiome’), we chose to use Faith’s phylogenetic diversity (47) (see Supplementary Text 1).

### Bayesian model fitting

We used a Bayesian framework to fit our basic model 1 (eq. 3) using Stan (48) and Rstan (49) to the gut and oral microbiome samples for the five separate groups: placebo, ciprofloxacin, clindamycin, minocycline, and amoxicillin (i.e. *n=*2×5=10 fits). In brief, our approach used 4 chains with a burn-in period of 1,000 iterations and 9,000 subsequent iterations, verifying that all chains converged 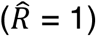 and the effective sample size for each parameter was sufficiently large (*n_eff* > 1,000). We additionally fitted model 2 with a possible state transition (eq. 4) to all non-placebo groups (*n=*2×4=8 fits).

We used non-informative priors for all parameters in the original model 1 without a state transition (eq. 3). For all groups, we used the same uniformly distributed prior for D (positive i.e. decrease in diversity) and uniform priors for *Φ*_5_, *Φ*_D_. For fitting model 2, we used an additional uniform prior centred at zero for the new equilibrium value *A* and the same priors for other parameters. In summary, the priors are as follows:

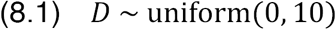

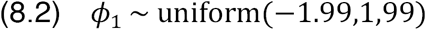

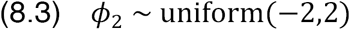

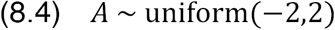

We compared models 1 and 2 (Supplementary Files 8 and 9). for each antibiotic treatment group using the Bayes factor (50,51) after extracting the model fits using bridge sampling with the bridgesampling R package v0.2-2 (52). A prior sensitivity analysis (not shown) showed that choice of priors did not affect our conclusions about model selection, although the strength of the Bayes factor varied.

Full code for fitting the models to empirical data and reproducing figures is available with this article (Supplementary Files 5—9).

### Shotgun metagenomics dataset

For fitting to the dataset released by Palleja et al. (27), we used the published species richness data with an arbitrary scaling factor (see Supplementary Text 1).

### Lotka-Volterra simulations

We numerically simulated 5^9^ = 1 953 125 parameter sets of the GLV model with *n=*3 species and investigated their behaviour and stable states. For more details see the corresponding supplementary discussion (Supplementary Text 2) and Mathematica notebook (Supplementary File 12).

