## Supplementary information - description of files for "Modelling microbiome recovery after antibiotics using a stability landscape framework"

### Supplemental Information

Figures

1. **Supplementary Figure 1 – Differences in individual response over time for the top twelve most abundant taxonomic families for placebo, clindamycin, and ciprofloxacin.** Relative abundances (log-scale) of the top twelve most abundant bacterial families plotted at each sampled timepoint. Observations are linked by coloured lines for each individual. Despite some consistency in changes between antibiotics across individuals, there is inter-individual variability and evidence of possible interactions between bacterial families.
2. **Supplementary Figure 2 – Personalised microbiome trajectories for each individual show a general structuring by antibiotic treatment group and body site.** Points represent the mean bootstrapped (n=100) phylogenetic diversity displacement relative to equilibrium for each individual in the dataset, with samples from the same individual connected by lines.
3. **Supplementary Figure 3 – Model fits to a rescaled dataset of species richness from Palleja et al. (2018).** See Supplementary Text 1 for details of reanalysis, and Supplementary File 6 for code.

Files

4. **Supplementary File 1 – Stan code defining Model 1 (no state transition).** For use with Supplementary Files 3 and 6.
5. **Supplementary File 2 – Stan code defining Model 2 (with state transition).** For use with Supplementary Files 3 and 6.
6. **Supplementary File 3 – All main analyses.** R markdown notebook for reproduction of the Zaura et al. analysis results in this paper, containing all analysis code. If run using Supplementary Files 1—5 this notebook produces: data files of bootstrapped phylogenetic diversity for all individuals; model fits; and resulting figures (Figures 2—4). A full archive including cached model fits and results is available on FigShare: <https://figshare.com/s/d62d6e90f96dc63c2769>.
7. **Supplementary File 4 – R phyloseq object containing reanalyzed gut microbiome data from Zaura et al. (2015).** For use with Supplementary File 3.
8. **Supplementary File 5 – R phyloseq object containing reanalyzed oral microbiome data from Zaura et al. (2015).** For use with Supplementary File 3.
9. **Supplementary File 6 – Palleja et al. (2018) analysis.** R code for fitting model to Palleja et al. (2018) dataset. See Supplementary Text 1 for details of reanalysis. For use with Supplementary Files 1, 2, and 7.
10. **Supplementary File 7 –** **Rarefied mOTU abundances from Palleja et al. (2018).** Original data from Palleja et al.’s analysis (downloaded from <http://arumugamlab.sund.ku.dk/SuppData/Palleja_et_al_2017_ABX/>) provided here to as a permanent link. For use with Supplementary File 6.
11. **Supplementary File 8 – Mathematica notebook of Lotka-Volterra simulations.** Interactive notebook containing code necessary to reproduce the analysis and figures in Supplementary Text 2.

Text

12. **Supplementary Text 1 – Additional details of methods and analysis.**
13. **Supplementary Text 2 – Details of Lotka-Volterra model simulations.** Detailed text and discussion reporting numerical simulations investigating behaviour predicted from the stability landscape framework using a Lotka-Volterra model in 3 dimensions.
