## Supplementary material for "Modelling microbiome recovery after antibiotics using a stability landscape framework": Supplementary-File-2.stan

```
functions {
  real model2( real t, real D, real phi1, real phi2, real Asym ){
    real y = D * exp(phi1) * exp(phi2) / (exp(phi2) - exp(phi1)) * (exp(-exp(phi1)*t) - exp(-exp(phi2)*t)) + Asym*(1-exp(-exp(phi1)*t));
    return y;
  }
}
data {
  int<lower=0> N;         // number of data points
  int<lower=0> n_subject;  // number of subjects
  vector[N] times;       // the times
  vector[N] concs;         // the concentrations
  int subjects[N];          // which subject, which time point
}

parameters {
  // for each parameter we have a mean value and a per-subject effect
  // as well as a parameter to represent the deviation of the treatment effects (used as s.d. in the normal distribution)
  // see pg. 208 of the STAN manual
  real<lower=0> D;
  real<lower=-2, upper=2> phi1; // log(0.5) to log(2)
  real<lower=-2, upper=2> phi2; // log(1) to log(4)
  real<lower=-2, upper=2> Asym;
  real<lower=0> sigma; // Standard deviation of individual observations
}

transformed parameters {
  real<lower=0> constr1;
  real<lower=0> constr2;
  constr1 = (exp(phi1) + exp(phi2))^2 - 4*exp(phi1)*exp(phi2); // so we are in overdamped parameter regime after transformation
  constr2 = phi2 - phi1; // restrict phi2>phi1
}

model {
  vector[N] fitted;

  // priors
  D ~ uniform(0, 10);
  phi1 ~ uniform(-1.99, 1.99);
  phi2 ~ uniform(-2, 2);
  Asym ~ uniform(-2, 2);
  // Asym ~ normal(0, 1.263);

  for (n in 1:N){
    fitted[n] = model2(times[n], D, phi1, phi2, Asym);

  }

  concs ~ normal(fitted, sigma);
}
```
