## Supplementary material for "Modelling microbiome recovery after antibiotics using a stability landscape framework": Supplementary-File-3.Rmd

````
---
title: "Analysis file: Modelling microbiome recovery after antibiotics using a stability landscape framework"
output: 
  html_document:
    number_sections: yes
    fig_width: 12
    fig_height: 10
    code_folding: hide
    results: hide

  cache: yes
    
author: "Liam P. Shaw,"

---

```{r load-libraries, results=FALSE, warnings=FALSE}
# Required libraries
require(phyloseq)
require(vegan)
require(picante)
suppressPackageStartupMessages(library(dplyr))
suppressPackageStartupMessages(library(tidyr))
suppressPackageStartupMessages(library(ggplot2))
library(rstan)
library(bridgesampling)
library(reshape2)
require(broman)
require(knitr)

cacheing=TRUE
```

# Introduction

This Rmarkdown file contains code to reproduce the main analyses of 

*Modelling microbiome recovery after antibiotics using a stability landscape framework*
  Liam P. Shaw, Hassan Bassam, Chris P. Barnes, A. Sarah Walker, Nigel Klein, Francois Balloux (2018).

The code in this file: calculates bootstrapped diversity displacements for each individual in the Zaura et al. (2015) study (SRA accession: SRP057504); fits models 1 and 2 to this data; and plots the resulting figures included in the associated main paper. Running all this analysis `from scratch' takes ~2 hours on a laptop with 16GB RAM. 

A previously cached version (i.e. pre-run and with filled folders) is available at Figshare: https://doi.org/10.6084/m9.figshare.6754880

Before running this markdown file, the directory structure should be constructed as follows, using the other supplementary files:

**`data`**

* `Supplementary-File-4.rds` -- `phyloseq` object of reanalyzed data from gut microbiome 
* `Supplementary-File-5.rds` -- `phyloseq` object of reanalyzed data from oral microbiome 

**`figures`**

empty (when run, will contain versions of figures from manuscript)

**`fitted-objects`**

empty (when run, contains objects storing fitted models)

**`models`**

* `Supplementary-File-1.stan` -- Stan model without alternative stable state parameter 
* `Supplementary-File-2.stan` -- Stan model with alternative stable state parameter


# Filtering and reanalysis of Zaura et al. (2015) data

The reanalysis of this data (not shown) proceeded as follows:

**Pre-processing**

* Download and combine all fastq files from original study using `fastq-dump`
* Filter based on maximum errors using `vsearch --fastq_filter all.fastq --fastq_maxee 1 --fastaout seqs.fa --fasta_width 0`
* Discard sequences with <395 bases (from inspection of length distribution of sequences) using `vsearch --fastq_filter all.fastq --fastq_minlen 395 --fastq_maxee 1 --fasta_width 0 --fastaout seqs.maxee_1.minlen_395.fa`
* Trim all sequences to 415 bases (from inspection of length distribution of sequences) using `awk '{print $1}' seqs.maxee_1.minlen_395.fa | sed -e 's/[.]/_/g' | cut -c 1-415 > seqs.maxee_1.minlen_395.trunc_415.fa`

**OTU clustering**

* Dereplication: `vsearch --derep_fulllength $seqs -output seqs.derep.fa -sizeout`
* Abundance sort and discard singletons: `vsearch -sortbysize seqs.derep.fa -output sorted.min2.fa -minsize 2`
* OTU clustering: `vsearch --cluster_fast sorted.min2.fa --usersort --id 0.97 --centroids cluster_fast.sorted.min2.centroids.fa; mv cluster_fast.sorted.min2.centroids.fa otus1.fa`
* De novo chimera filtering: `vsearch --uchime_denovo otus1.fa --nonchimeras otus.nonchimeras.de.novo.fa -strand plus`
* Chimera filtering using 'gold' reference database (http://drive5.com/uchime/gold.fa): `vsearch -uchime_ref otus.nonchimeras.de.novo.fa -db gold.fa -strand plus -nonchimeras otus.nonchimeras.de.novo.gold.fa`
* Map original reads back to OTUs: `vsearch -usearch_global $seqs -db otus.nonchimeras.de.novo.gold.fa -strand plus -id 0.97 -uc otu_map.uc`
* Assign taxonomy to the OTUs: `parallel_assign_taxonomy_rdp.py -r /Users/liam/gg_13_5_otus/rep_set/99_otus.fasta -t /Users/liam/gg_13_5_otus/taxonomy/99_otu_taxonomy.txt -i otus.nonchimeras.de.novo.gold.fa -o rdp-assigned-tax/ -O 4 -v`
* Run FastTree to get marker gene phylogeny: `FastTree -nt -gtr otus.nonchimeras.de.novo.gold.fa > otus.nonchimeras.de.novo.gold.fa.FastTree.tre`


# Bootstrapping to obtain phylogenetic diversity

We first load the prepared files for the gut and oral microbiome datasets respectively, then calculate bootstrapped mean phylogenetic diversity displacements from baseline for each individual. 

```{r bootstrap-diversity-displacement-functions, cache=cacheing, warning=FALSE}
# Function to calculate data for a single participant in the study
participantData <- function(participant, a.phyloseq, N=100, rarefy.depth=1000){
  participant.data <- prune_samples(a.phyloseq, samples=sample_data(a.phyloseq)$participant==participant)
  example.rarefaction <-   rarefy_even_depth(participant.data, sample.size=rarefy.depth)
  # Get rid of samples with insufficient samples
  participant.data <- prune_samples(participant.data, samples=sample_names(example.rarefaction))
  participant.values <- matrix(nrow=N, ncol=nsamples(participant.data))
  # Get order of timepoints (not always as expected)
  sample.order <- as.numeric(sample_data(example.rarefaction)$time_s) 
  # Iterate through and bootstrap phylogenetic diversity
  for (i in 1:N){
    #print(i)
    participant.data.rarefy <- rarefy_even_depth(participant.data, sample.size=rarefy.depth)
    sample <- data.frame(t(otu_table(participant.data.rarefy)), check.names=F)
    faith.pd <- pd(sample, tree=prune.sample(sample, phy_tree(participant.data.rarefy)), include.root=FALSE)
    participant.values[i,] <- faith.pd$PD[sample.order] # order it!
  }
  # Subtract initial values to obtain diversity displacements
  participant.values <- -data.frame(participant.values-participant.values[,1])
  # Preparing data frame
  colnames(participant.values) <- c("0", "11", "30", "60", "120", "365")
  participant.values$rarefaction <- seq(1,N)
  empirical.data <- melt(participant.values, id.vars=c("rarefaction"))
  colnames(empirical.data) <- c("rarefaction", "t", "y")
  empirical.data$t <- as.numeric(as.character(empirical.data$t))
  return(empirical.data)
}

# Function to calculate bootstrapped diversity displacements for all samples in a dataset
allBootstrappedData <- function(a.phyloseq, csv.filename, bootstraps=100, rarefy.depth=1000){
  # Check rarefaction
  rarefied <- rarefy_even_depth(a.phyloseq, sample.size=rarefy.depth)
  s <- data.frame(sample_data(rarefied))
  participant.names <- unique(s$participant)
  six.obs <- table(s$participant)==6 # only keep those with 6 observations
  participant.names <- participant.names[six.obs[participant.names]=="TRUE"]
  
  
  for (p in participant.names){
    print(p)
    data <- participantData(p, a.phyloseq, N = bootstraps, rarefy.depth = rarefy.depth)
    means <- as.data.frame(data %>% group_by(t) %>% summarise_all(mean))
    sds <- as.data.frame(data %>% group_by(t) %>% summarise_all(sd))
    
    means$rarefaction <- NULL
    means$sd <- sds$y
    means$participant <- p
    means$antibiotic <- unique(s[s$participant==p, "chem_administration_s"])
    means$centre <- unique(s[s$participant==p, "centre_s"])
    
    # Append data to file
    write.table(means, file=csv.filename, append = TRUE, sep = ",", row.names = F, col.names = F)
  }
}
```
```{r bootstrap-diversity-oral, cache=cacheing, warning=FALSE}
###################
# Oral microbiome #
###################
# Uncomment to run bootstrapping
## Read in data
oral <- readRDS('data/Supplementary-File-5.rds')
# Calculate bootstrapped diversity displacements
allBootstrappedData(oral, csv.filename = 'data/oral_PD_100_mean.csv', bootstraps=100)
```

```{r bootstrap-diversity-gut, cache=cacheing, warning=FALSE}
###################
# Gut microbiome #
###################
# Uncomment to run bootstrapping 
## Read in data
 gut <- readRDS('data/Supplementary-File-4.rds')
## Calculate bootstrapped diversity displacements
 allBootstrappedData(gut, csv.filename = 'data/gut_PD_100_mean.csv', bootstraps=100)
```

# Fitting models with Stan

We use Stan to fit our model to the experimental data.

```{r fitmodels, cache=cacheing, warning=FALSE, results=FALSE}
# Default theme
theme_set(theme_bw() + theme(panel.margin = grid::unit(0, "lines")))

set.seed(4711)

### USEFUL FUNCTIONS ###
# The original model (model 1)
ssfol <- function(D, phi1, phi2, t){
  y <- D * exp(phi1) * exp(phi2) / (exp(phi2) - exp(phi1)) * 
    (exp(-exp(phi1)*t) - exp(-exp(phi2)*t))
  return(y)
}
# The model plus asymptote (model 2)
ssfolAsym <- function(D, phi1, phi2, Asym, t){
  y <- D * exp(phi1) * exp(phi2) / (exp(phi2) - exp(phi1)) * 
    (exp(-exp(phi1)*t) - exp(-exp(phi2)*t)) + Asym*(1-exp(-exp(phi1)*t))
  return(y)
}
# Compare two models
compare_models <- function(model1, model2, method="normal"){
  bs.model1 <- bridge_sampler(model1, method = method)
  bs.model2 <- bridge_sampler(model2, method = method)
  return(bf(bs.model1, bs.model2))
}

# STAN Models
## MODELS TO USE
####
# Initial model (no alternative stable state)
model.no.asym <- 'models/Supplementary-File-1.stan' 
# Model 2 (alternative stable state allowed)
model <- 'models/Supplementary-File-2.stan' 

# PARAMETERS
iterations <- 10000
warmups <- 1000

# Function to fit stan models and calculate Bayes factor
fitStan <- function(antibiotic, data, site){
  antibiotic.df <- data[data$V5==antibiotic,]
  antibiotic.data <- list(N=nrow(antibiotic.df), n_subject=nrow(antibiotic.df)/6, concs=antibiotic.df$V2, times=antibiotic.df$V1/30, subjects=as.vector(sapply(seq(1, nrow(antibiotic.df)/6), function(x) rep(x, 6))))
  antibiotic.data <- list(N=nrow(antibiotic.df), n_subject=nrow(antibiotic.df)/6, concs=antibiotic.df$V2, times=antibiotic.df$V1/30, subjects=as.vector(sapply(seq(1, nrow(antibiotic.df)/6), function(x) rep(x, 6))))
  # Fit model 1 (without asymptote)
  mr.stan.no.asym =  stan(model.no.asym, chains = 4, iter = iterations, data = antibiotic.data,warmup = warmups)
  # Fit model 2 (with asymptote)
  mr.stan =  stan(model, chains = 4, iter = iterations, data = antibiotic.data, warmup = warmups)
  # Compare the two models
  bayes.factor <- compare_models(mr.stan, mr.stan.no.asym)
  # Save objects
  saveRDS(antibiotic.data, file=paste('fitted-objects/', site, '-', antibiotic, '-points.rds', sep=''))
  saveRDS(bayes.factor, file=paste('fitted-objects/', site, '-', antibiotic, '-bayes-factor.rds', sep=''))
  saveRDS(mr.stan, file=paste('fitted-objects/', site, '-', antibiotic, '-full-model.rds', sep=''))
  saveRDS(mr.stan.no.asym, file=paste('fitted-objects/', site, '-', antibiotic, '-no-asym-model.rds', sep=''))
}

###################
# Oral microbiome #
##################

# Read in precalculated bootstrapped data 
oral <- read.csv('data/oral_PD_100_mean.csv', header=F)

# Four antibiotics
fitStan("clinda", oral, "oral")
fitStan("cipro", oral, "oral")
fitStan("minoc", oral, "oral")
fitStan("amox", oral, "oral")

# Placebo - only fit model without asymptote
placebo <- oral[oral$V5=="placebo",]
placebo.data <- list(N=nrow(placebo), n_subject=nrow(placebo)/6, concs=placebo$V2, times=placebo$V1/30, subjects=as.vector(sapply(seq(1, nrow(placebo)/6), function(x) rep(x, 6))))
saveRDS(placebo.data, file='fitted-objects/oral-placebo-points.rds')
# Fit
mr.stan.no.asym =  stan(model.no.asym, chains = 4, iter = iterations, data = placebo.data,warmup = warmups)
# Save
saveRDS(mr.stan.no.asym, file=paste('fitted-objects/oral-', 'placebo', '-no-asym-model.rds', sep=''))

##################
# Gut microbiome #
##################

# Read in precalculated data 
gut <- read.csv('data/gut_PD_100_mean.csv', header=F)
fitStan("clinda", data=gut, site='gut')
fitStan("cipro", data=gut, site='gut')
fitStan("minoc", data=gut, site='gut')
fitStan("amox", data=gut, site='gut')

clinda.bf <- readRDS('fitted-objects/gut-clinda-bayes-factor.rds')
cipro.bf <- readRDS('fitted-objects/gut-cipro-bayes-factor.rds')
minoc.bf <- readRDS('fitted-objects/gut-minoc-bayes-factor.rds')
amox.bf <- readRDS('fitted-objects/gut-amox-bayes-factor.rds')

# Placebo - only fit model without asymptote
placebo <- gut[gut$V5=="placebo",]
placebo.data <- list(N=nrow(placebo), n_subject=nrow(placebo)/6, concs=placebo$V2, times=placebo$V1/30, subjects=as.vector(sapply(seq(1, nrow(placebo)/6), function(x) rep(x, 6))))
saveRDS(placebo.data, file='fitted-objects/gut-placebo-points.rds')
# Fit
mr.stan.no.asym =  stan(model.no.asym, chains = 4, iter = iterations, data = placebo.data,warmup = warmups)
# Save
saveRDS(mr.stan.no.asym, file=paste('fitted-objects/gut-', 'placebo', '-no-asym-model.rds', sep=''))
```

# Plotting 

Now we have everything we need to produce versions of the final figures in the paper. 

```{r plot-fits, cache=cacheing}
# Multiplot function from http://www.cookbook-r.com/Graphs/Multiple_graphs_on_one_page_(ggplot2)/
multiplot <- function(..., plotlist=NULL, file, cols=1, layout=NULL) {
  library(grid)
  
  # Make a list from the ... arguments and plotlist
  plots <- c(list(...), plotlist)
  
  numPlots = length(plots)
  
  # If layout is NULL, then use 'cols' to determine layout
  if (is.null(layout)) {
    # Make the panel
    # ncol: Number of columns of plots
    # nrow: Number of rows needed, calculated from # of cols
    layout <- matrix(seq(1, cols * ceiling(numPlots/cols)),
                     ncol = cols, nrow = ceiling(numPlots/cols))
  }
  
  if (numPlots==1) {
    print(plots[[1]])
    
  } else {
    # Set up the page
    grid.newpage()
    pushViewport(viewport(layout = grid.layout(nrow(layout), ncol(layout))))
    
    # Make each plot, in the correct location
    for (i in 1:numPlots) {
      # Get the i,j matrix positions of the regions that contain this subplot
      matchidx <- as.data.frame(which(layout == i, arr.ind = TRUE))
      
      print(plots[[i]], vp = viewport(layout.pos.row = matchidx$row,
                                      layout.pos.col = matchidx$col))
    }
  }
}

# Function to make plot of model and points
makePlot <- function(data.df, stan.model, asymptote=TRUE, model.colour='grey', ylimits=c(-5, 8), title=""){
  plot(data.df$times, data.df$concs, pch=19, xlab="", ylab="", ylim=ylimits,
       cex.lab=2, cex.axis=2)
  times <- seq(0, 12, 0.1)
  stan.model.ex <- extract(stan.model)
  points <- matrix(nrow=100000, ncol=length(times))
  for(i in seq(1,100000)){
    if (asymptote==TRUE){
      points[i,] <- ssfolAsym(D=stan.model.ex$D[i], phi1=stan.model.ex$phi1[i], 
                              phi2=stan.model.ex$phi2[i], Asym=stan.model.ex$Asym[i], times)
    }
    else{
      points[i,] <- ssfol(D=stan.model.ex$D[i], phi1=stan.model.ex$phi1[i], 
                          phi2=stan.model.ex$phi2[i],times)
    }
  }
  # Add zero line
  abline(a=0,b=0, lwd=3, lty=2, col='grey')
  
  
  # Add 2.5%, median, 97.5% lines
  points.lower <-  apply(na.omit(points), function(x) quantile(x, probs = 0.025), MARGIN=2)
  points.median <- apply(na.omit(points), function(x) quantile(x, probs = 0.5), MARGIN=2)
  points.upper <- apply(na.omit(points), function(x) quantile(x, probs=0.975), MARGIN=2)
  points(times, points.median, type='l', col=model.colour, lwd=2)
  points(times, points.lower, type='l', col=model.colour, lwd=2, lty=2)
  points(times, points.upper, type='l', col=model.colour, lwd=2, lty=2)
  
  # add data points on top
  points(data.df$times, data.df$concs, pch=19)
  
}

# Function for model with/without alternative stable state
makeAntibioticPlotNoAsym <- function(antibiotic, site, model.colour){
  antibiotic.data <- readRDS(paste('fitted-objects/', site, '-', antibiotic, '-points.rds', sep=''))
  model <- readRDS(paste('fitted-objects/', site, '-', antibiotic, '-no-asym-model.rds', sep=''))
  makePlot(antibiotic.data, model, asymptote=FALSE, model.colour=model.colour)
}
makeAntibioticPlotAsym <- function(antibiotic, site, model.colour){
  antibiotic.data <- readRDS(paste('fitted-objects/', site, '-', antibiotic, '-points.rds', sep=''))
  model <- readRDS(paste('fitted-objects/', site, '-', antibiotic, '-full-model.rds', sep=''))
  makePlot(antibiotic.data, model, asymptote=TRUE, model.colour=model.colour)
}
```

## Figure 2

See `figures/Figure-2-model-1-fits.pdf`

```{r figure2, warning=FALSE, cache=cacheing}
############
# FIGURE 2 #
############
# Model 1 fits (corresponding to Figure 2 of paper)
pdf('figures/Figure-2-model-1-fits.pdf', width=25, height=10)
par(mfrow=c(2,5))
# Gut microbiome
makeAntibioticPlotNoAsym("placebo", "gut", "blue")
makeAntibioticPlotNoAsym("cipro", "gut", "green")
makeAntibioticPlotNoAsym("clinda", "gut", "red")
makeAntibioticPlotNoAsym("minoc", "gut", "purple")
makeAntibioticPlotNoAsym("amox", "gut", "orange")
# Oral microbiome
makeAntibioticPlotNoAsym("placebo", "oral", "blue")
makeAntibioticPlotNoAsym("cipro", "oral", "green")
makeAntibioticPlotNoAsym("clinda", "oral", "red")
makeAntibioticPlotNoAsym("minoc", "oral", "purple")
makeAntibioticPlotNoAsym("amox", "oral", "orange")
dev.off()
```

## Figure 3

See `figures/Figure-3-model-2-fits.pdf`

```{r figure3, warning=FALSE, cache=cacheing}
############
# FIGURE 3 #
############
# Model 2 fits (corresponding to Figure 3 of paper)
pdf('figures/Figure-3-model-2-fits.pdf', width=20, height=10)
par(mfrow=c(2,4))
# Gut microbiome
makeAntibioticPlotAsym("cipro", "gut", "green")
makeAntibioticPlotAsym("clinda", "gut", "red")
makeAntibioticPlotAsym("minoc", "gut", "purple")
makeAntibioticPlotAsym("amox", "gut", "orange")
# Oral microbiome
makeAntibioticPlotAsym("cipro", "oral", "green")
makeAntibioticPlotAsym("clinda", "oral", "red")
makeAntibioticPlotAsym("minoc", "oral", "purple")
makeAntibioticPlotAsym("amox", "oral", "orange")
dev.off()
```

## Figure 4

See `figures/Figure-4-parameter-plots.pdf`.
```{r figure4, warning=FALSE, cache=cacheing}
############
# FIGURE 4 #
############
## PARAMETER PLOTS
# Read in data
cipro.oral <- rstan::extract(readRDS('fitted-objects/oral-cipro-full-model.rds'))
clinda.oral <- rstan::extract(readRDS('fitted-objects/oral-clinda-full-model.rds'))
minoc.oral <- rstan::extract(readRDS('fitted-objects/oral-minoc-full-model.rds'))
amox.oral <- rstan::extract(readRDS('fitted-objects/oral-amox-full-model.rds'))

cipro.gut <- rstan::extract(readRDS('fitted-objects/gut-cipro-full-model.rds'))
clinda.gut <- rstan::extract(readRDS('fitted-objects/gut-clinda-full-model.rds'))
minoc.gut <- rstan::extract(readRDS('fitted-objects/gut-minoc-full-model.rds'))
amox.gut <- rstan::extract(readRDS('fitted-objects/gut-amox-full-model.rds'))


# PARAMETER PLOTTING
ylimits <- c(0, 1.75)
# Parameter: D
sample.size <- 10000
d.oral <- cbind(sample(na.omit(cipro.oral$D), size = sample.size),
                sample(na.omit(clinda.oral$D), size = sample.size),
                sample(na.omit(minoc.oral$D), size = sample.size),
                sample(na.omit(amox.oral$D), size = sample.size))
colnames(d.oral) <- c("Ciprofloxacin", "Clindamycin", "Minocycline", "Amoxicillin")
d.oral.melt <- melt(d.oral)
d.gut <- cbind(sample(na.omit(cipro.gut$D), size = sample.size),
                sample(na.omit(clinda.gut$D), size = sample.size),
                sample(na.omit(minoc.gut$D), size = sample.size),
                sample(na.omit(amox.gut$D), size = sample.size))
colnames(d.gut) <- c("Ciprofloxacin", "Clindamycin", "Minocycline", "Amoxicillin")
d.gut.melt <- melt(d.gut)
p.d <- ggplot(d.oral.melt, aes(x=value, group=Var2, colour=Var2))+
  geom_density(linetype='dashed')+
  scale_color_manual(values=c("green", "red", "purple", "orange"))+
  geom_density(data=d.gut.melt, aes(x=value, group=Var2, colour=Var2))+
  facet_wrap(~Var2, ncol=1)+
  theme(axis.text=element_text(colour='black'))+
  theme(legend.position="none")+
  ylim(c(0,0.75))+
  xlim(c(0,10))+
  xlab("")+ylab("")+theme(strip.background=element_blank(), strip.text=element_blank())+
    theme_bw()+
    guides(colour=FALSE)+
  ggtitle("Strength of perturbation (D)")+
  theme(plot.title = element_text(size=18, hjust=0.5))

# Parameter: asymptote
asym.oral <- cbind(sample(na.omit(cipro.oral$Asym), size = sample.size),
                sample(na.omit(clinda.oral$Asym), size = sample.size),
                sample(na.omit(minoc.oral$Asym), size = sample.size),
                sample(na.omit(amox.oral$Asym), size = sample.size))
colnames(asym.oral) <- c("Ciprofloxacin", "Clindamycin", "Minocycline", "Amoxicillin")
asym.oral.melt <- melt(asym.oral)
asym.gut <- cbind(sample(na.omit(cipro.gut$Asym), size = sample.size),
               sample(na.omit(clinda.gut$Asym), size = sample.size),
               sample(na.omit(minoc.gut$Asym), size = sample.size),
               sample(na.omit(amox.gut$Asym), size = sample.size))
colnames(asym.gut) <- c("Ciprofloxacin", "Clindamycin", "Minocycline", "Amoxicillin")
asym.gut.melt <- melt(asym.gut)
p.asym <- ggplot(asym.oral.melt, aes(x=value, group=Var2, colour=Var2))+
  geom_density(linetype='dashed')+
  scale_color_manual(values=c("green", "red", "purple", "orange"))+
  geom_density(data=asym.gut.melt, aes(x=value, group=Var2, colour=Var2))+
  facet_wrap(~Var2, ncol=1)+
  theme(axis.text=element_text(colour='black'))+
  theme(legend.position="none")+
  ylim(c(0,1.5))+
  xlab("")+ylab("")+theme(strip.background=element_blank(), strip.text=element_blank())+
    theme_bw()+
    guides(colour=FALSE)+
  ggtitle("Asymptote parameter (A)")+
  theme(plot.title = element_text(size=18, hjust=0.5))

# PARAMETER: phi1
phi1.oral <- cbind(sample(na.omit(cipro.oral$phi1), size = sample.size),
                   sample(na.omit(clinda.oral$phi1), size = sample.size),
                   sample(na.omit(minoc.oral$phi1), size = sample.size),
                   sample(na.omit(amox.oral$phi1), size = sample.size))
colnames(phi1.oral) <- c("Ciprofloxacin", "Clindamycin", "Minocycline", "Amoxicillin")
phi1.oral.melt <- melt(phi1.oral)
phi1.gut <- cbind(sample(na.omit(cipro.gut$phi1), size = sample.size),
                  sample(na.omit(clinda.gut$phi1), size = sample.size),
                  sample(na.omit(minoc.gut$phi1), size = sample.size),
                  sample(na.omit(amox.gut$phi1), size = sample.size))
colnames(phi1.gut) <- c("Ciprofloxacin", "Clindamycin", "Minocycline", "Amoxicillin")
phi1.gut.melt <- melt(phi1.gut)
p.phi1 <- ggplot(phi1.oral.melt, aes(x=value, group=Var2, colour=Var2))+
  geom_density(linetype='dashed')+
  scale_color_manual(values=c("green", "red", "purple", "orange"))+
  geom_density(data=phi1.gut.melt, aes(x=value, group=Var2, colour=Var2))+
  facet_wrap(~Var2, ncol=1)+
  theme(axis.text=element_text(colour='black'))+
  theme(legend.position="none")+
  ylim(ylimits)+
  xlab("")+ylab("")+theme(strip.background=element_blank(), strip.text=element_blank())


# Parameter: phi2
phi2.oral <- cbind(sample(na.omit(cipro.oral$phi2), size = sample.size),
                   sample(na.omit(clinda.oral$phi2), size = sample.size),
                   sample(na.omit(minoc.oral$phi2), size = sample.size),
                   sample(na.omit(amox.oral$phi2), size = sample.size))
colnames(phi2.oral) <- c("Ciprofloxacin", "Clindamycin", "Minocycline", "Amoxicillin")
phi2.oral.melt <- melt(phi2.oral)
phi2.gut <- cbind(sample(na.omit(cipro.gut$phi2), size = sample.size),
                  sample(na.omit(clinda.gut$phi2), size = sample.size),
                  sample(na.omit(minoc.gut$phi2), size = sample.size),
                  sample(na.omit(amox.gut$phi2), size = sample.size))
colnames(phi2.gut) <- c("Ciprofloxacin", "Clindamycin", "Minocycline", "Amoxicillin")
phi2.gut.melt <- melt(phi2.gut)
p.phi2 <- ggplot(phi2.oral.melt, aes(x=value, group=Var2, colour=Var2))+
  geom_density(linetype='dashed')+
  scale_color_manual(values=c("green", "red", "purple", "orange"))+
  geom_density(data=phi2.gut.melt, aes(x=value, group=Var2, colour=Var2))+
  facet_wrap(~Var2, ncol=1)+
  theme(axis.text=element_text(colour='black'))+
  theme(legend.position="none")+
  ylim(ylimits)+
  xlab("")+ylab("")+theme(strip.background=element_blank(), strip.text=element_blank())

# PARAMETER: b
ylimits.b.k <-  c(0,3.75)
b.oral.melt <- melt(exp(phi1.oral) + exp(phi2.oral))
b.gut.melt <- melt(exp(phi1.gut) + exp(phi2.gut))
p.b <- ggplot(b.oral.melt, aes(x=value, group=Var2, colour=Var2))+
  geom_density(linetype='dashed')+
  scale_color_manual(values=c("green", "red", "purple", "orange"))+
  geom_density(data=b.gut.melt, aes(x=value, group=Var2, colour=Var2))+
  facet_wrap(~Var2, ncol=1)+
  theme(axis.text=element_text(colour='black'))+
  theme(legend.position="none")+
  #ylim(ylimits)+
  xlab("")+ylab("")+theme(strip.background=element_blank(), strip.text=element_blank())+
  xlim(c(0,10))+
  ylim(c(0,1))+
  theme(axis.text.y = element_blank())+
    theme_bw()+
    guides(colour=FALSE)+
  ggtitle("Strength of damping (b)")+
  theme(plot.title = element_text(size=18, hjust=0.5))

# PARAMETER: k
k.oral.melt <- melt(exp(phi1.oral + phi2.oral))
k.gut.melt <- melt(exp(phi1.gut + phi2.gut))
p.k <- ggplot(k.oral.melt, aes(x=value, group=Var2, colour=Var2))+
  geom_density(linetype='dashed')+
  scale_color_manual(values=c("green", "red", "purple", "orange"))+
  geom_density(data=k.gut.melt, aes(x=value, group=Var2, colour=Var2))+
  facet_wrap(~Var2, ncol=1)+
  theme(axis.text=element_text(colour='black'))+
  theme(legend.position="none")+
  #ylim(ylimits)+
  xlab("")+ylab("")+theme(strip.background=element_blank(), strip.text=element_blank())+
  xlim(c(0,5))+
  ylim(ylimits.b.k)+
  theme(axis.text.y = element_blank())+
  theme_bw()+
  guides(colour=FALSE)+
  ggtitle("Strength of restoring force (k)")+
  theme(plot.title = element_text(size=18, hjust=0.5))

  # Multiplot
library(easyGgplot2)
pdf('figures/Figure-4-parameter-plots.pdf', width=15, height=10)
easyGgplot2::ggplot2.multiplot(p.d, p.asym, p.b, p.k, cols = 4)
dev.off()
```


## Table 3

We save the 95% credible intervals for all parameters for the model fits as separate tables, which are then combined to produce Table 3 in the main manuscript. Please note that due to the stochastic nature of the fitting of the model there will likely be some small discrepancies in the intervals produced from this script compared to the published version. 

```{r supplementary-table-1, warning=FALSE, cache=cacheing}
# Gut
cipro.gut <- extract(readRDS('fitted-objects/gut-cipro-full-model.rds'))
clinda.gut <- extract(readRDS('fitted-objects/gut-clinda-full-model.rds'))
amox.gut <- extract(readRDS('fitted-objects/gut-amox-full-model.rds'))
minoc.gut <- extract(readRDS('fitted-objects/gut-minoc-full-model.rds'))
# Oral
cipro.oral <- extract(readRDS('fitted-objects/oral-cipro-full-model.rds'))
clinda.oral <- extract(readRDS('fitted-objects/oral-clinda-full-model.rds'))
amox.oral <- extract(readRDS('fitted-objects/oral-amox-full-model.rds'))
minoc.oral <- extract(readRDS('fitted-objects/oral-minoc-full-model.rds'))

# Make tables
rows <- c("cipro.gut", "clinda.gut", "amox.gut", "minoc.gut",
          "cipro.oral", "clinda.oral", "amox.oral", "minoc.oral")
D.posterior <- data.frame(rbind(myround(quantile(cipro.gut$D, probs = c(0.5, 0.05, 0.95)), 2),
      myround(quantile(clinda.gut$D, probs = c(0.5, 0.05, 0.95)), 2),
      myround(quantile(amox.gut$D, probs = c(0.5, 0.05, 0.95)), 2),
      myround(quantile(minoc.gut$D, probs = c(0.5, 0.05, 0.95)), 2),
      myround(quantile(cipro.oral$D, probs = c(0.5, 0.05, 0.95)), 2),
      myround(quantile(clinda.oral$D, probs = c(0.5, 0.05, 0.95)), 2),
      myround(quantile(amox.oral$D, probs = c(0.5, 0.05, 0.95)), 2),
      myround(quantile(minoc.oral$D, probs = c(0.5, 0.05, 0.95)), 2)))
colnames(D.posterior) <- c("median", "2", "3")
rownames(D.posterior) <- rows
D.posterior$credible.interval.95 <-  paste("(", D.posterior[,2], "--", D.posterior[,3], ")", sep="")
D.posterior[,c("2", "3")] <- NULL
write.csv(D.posterior, file='fitted-objects/parameter-credible-intervals-D.csv', row.names=TRUE, quote=T)
kable(D.posterior)

# Asym
Asym.posterior <- data.frame(rbind(myround(quantile(cipro.gut$Asym, probs = c(0.5, 0.05, 0.95)), 2),
      myround(quantile(clinda.gut$Asym, probs = c(0.5, 0.05, 0.95)), 2),
      myround(quantile(amox.gut$Asym, probs = c(0.5, 0.05, 0.95)), 2),
      myround(quantile(minoc.gut$Asym, probs = c(0.5, 0.05, 0.95)), 2),
      myround(quantile(cipro.oral$Asym, probs = c(0.5, 0.05, 0.95)), 2),
      myround(quantile(clinda.oral$Asym, probs = c(0.5, 0.05, 0.95)), 2),
      myround(quantile(amox.oral$Asym, probs = c(0.5, 0.05, 0.95)), 2),
      myround(quantile(minoc.oral$Asym, probs = c(0.5, 0.05, 0.95)), 2)))
colnames(Asym.posterior) <- c("median", "2", "3")
rownames(Asym.posterior) <- rows
Asym.posterior$credible.interval.95 <-  paste("(", Asym.posterior[,2], "--", Asym.posterior[,3], ")", sep="")
Asym.posterior[,c("2", "3")] <- NULL
write.csv(Asym.posterior, file='fitted-objects/parameter-credible-intervals-Asym.csv', row.names=TRUE, quote=T)
kable(Asym.posterior)

# Phi1
phi1.posterior <- data.frame(rbind(myround(quantile(cipro.gut$phi1, probs = c(0.5, 0.05, 0.95)), 2),
      myround(quantile(clinda.gut$phi1, probs = c(0.5, 0.05, 0.95)), 2),
      myround(quantile(amox.gut$phi1, probs = c(0.5, 0.05, 0.95)), 2),
      myround(quantile(minoc.gut$phi1, probs = c(0.5, 0.05, 0.95)), 2),
      myround(quantile(cipro.oral$phi1, probs = c(0.5, 0.05, 0.95)), 2),
      myround(quantile(clinda.oral$phi1, probs = c(0.5, 0.05, 0.95)), 2),
      myround(quantile(amox.oral$phi1, probs = c(0.5, 0.05, 0.95)), 2),
      myround(quantile(minoc.oral$phi1, probs = c(0.5, 0.05, 0.95)), 2)))
colnames(phi1.posterior) <- c("median", "2", "3")
rownames(phi1.posterior) <- rows
phi1.posterior$credible.interval.95 <- paste("(", phi1.posterior[,2], "--", phi1.posterior[,3], ")", sep="")
phi1.posterior[,c("2", "3")] <- NULL
write.csv(phi1.posterior, file='fitted-objects/parameter-credible-intervals-phi1.csv', row.names=TRUE, quote=T)
kable(phi1.posterior)

# Phi2
phi2.posterior <- data.frame(rbind(myround(quantile(cipro.gut$phi2, probs = c(0.5, 0.05, 0.95)), 2),
      myround(quantile(clinda.gut$phi2, probs = c(0.5, 0.05, 0.95)), 2),
      myround(quantile(amox.gut$phi2, probs = c(0.5, 0.05, 0.95)), 2),
      myround(quantile(minoc.gut$phi2, probs = c(0.5, 0.05, 0.95)), 2),
      myround(quantile(cipro.oral$phi2, probs = c(0.5, 0.05, 0.95)), 2),
      myround(quantile(clinda.oral$phi2, probs = c(0.5, 0.05, 0.95)), 2),
      myround(quantile(amox.oral$phi2, probs = c(0.5, 0.05, 0.95)), 2),
      myround(quantile(minoc.oral$phi2, probs = c(0.5, 0.05, 0.95)), 2)))
colnames(phi2.posterior) <- c("median", "2", "3")
rownames(phi2.posterior) <- rows
phi2.posterior$credible.interval.95 <- paste("(", phi2.posterior[,2], "--", phi2.posterior[,3], ")", sep="")
phi2.posterior[,c("2", "3")] <- NULL
write.csv(phi2.posterior, file='fitted-objects/parameter-credible-intervals-phi2.csv', row.names=TRUE, quote=T)
kable(phi2.posterior)
```
````
