## Supplementary material for "Modelling microbiome recovery after antibiotics using a stability landscape framework": Supplementary-File-6.R

```
# Reanalysis of data from
# Palleja et al. (2018)
# https://doi.org/10.1038/s41564-018-0257-9

# To be used in conjunction with Supplementary Files 1, 2, and 7 from
# the associated publication Shaw et al. (2019)

# Read in rarefied relative abundances
# N.B. Originally downloaded from http://arumugamlab.sund.ku.dk/SuppData/Palleja_et_al_2017_ABX/
t <- read.table('Supplementary-File-7.tsv', sep='\t', header=T, row.names = 1)


require(vegan)

shannon <- diversity(t(t), index = "shannon")
require(Matrix)
richness <- apply(t, FUN=nnzero, MARGIN=2)
df <- data.frame(cbind(shannon, richness, gsub(".*_", "", colnames(t))), gsub("_.*", "", colnames(t)))
colnames(df) <- c("shannon", "richness", "sample", "patient")
# Order samples
df$sample[which(df$sample=="Dag4opt")] <- "Dag4"
df$sample[which(df$sample=="Dag8opt")] <- "Dag8"
df$sample <- ordered(df$sample, levels=c("Dag0", "Dag4", "Dag8","Dag42", "Dag180"))

# Convert diversities to numeric
df$shannon <- as.numeric(as.character(df$shannon))
df$richness <- as.numeric(as.character(df$richness))

# Order by sample
df <- df[order(df$sample),]

# Normalised richness (relative to each individual baseline)
df$richness.norm <- sapply(rownames(df), 
                           function(x) df[x,"richness"] - df[which(df$patient==gsub("_.*", "", x) & df$sample=="Dag0"),"richness"])


# Date as actual number
df$day <- as.numeric(gsub("Dag", "", df$sample))

# Only keep those with all data points
patient.counts <- table(df$patient)
df.full.data <- names(patient.counts[which(patient.counts==5)])
df.write <- df[which(df$patient %in% df.full.data),c("day", "richness.norm")]
colnames(df.write) <- c("t", "y")
df.write$y <- -df.write$y/50 # arbitrary scaling for fitting
df.write$t <- df.write$t/30 # Convert day to month

# Write data to file
write.csv(df.write, file='rarefied-richness-relative.csv', quote=F, row.names = F)


# Fit

library(dplyr)
library(tidyr)
library(ggplot2)
library(rstan)
library(bridgesampling)
library(reshape2)
cat("libraries loaded")

set.seed(4711)

### USEFUL FUNCTIONS ###
# The original model (model 1)
ssfol <- function(D, phi1, phi2, t){
  y <- D * exp(phi1) * exp(phi2) / (exp(phi2) - exp(phi1)) * 
    (exp(-exp(phi1)*t) - exp(-exp(phi2)*t))
  return(y)
}
# The model plus asymptote (model 2)
ssfolAsym <- function(D, phi1, phi2, Asym, t){
  y <- D * exp(phi1) * exp(phi2) / (exp(phi2) - exp(phi1)) * 
    (exp(-exp(phi1)*t) - exp(-exp(phi2)*t)) + Asym*(1-exp(-exp(phi1)*t))
  return(y)
}
# Compare two models
compare_models <- function(model1, model2, method="normal"){
  bs.model1 <- bridge_sampler(model1, method = method)
  bs.model2 <- bridge_sampler(model2, method = method)
  return(bf(bs.model1, bs.model2))
}

# Change this to base directory of unzipped archive

## MODELS TO USE
model <- 'Supplementary-File-1.stan'
model.no.asym <- 'Supplementary-File-2.stan'

### LOAD DATA ####
n_subject <- 9
data.palleja <- read.csv('rarefied-richness-relative.csv', header=T)
palleja.data <- list(N=45, 
                     n_subject=9,
                     concs=data.palleja$y,
                     times=data.palleja$t, 
                     subjects=as.vector(rep(seq(1, 9), 5)))
### STAN MODEL FITTING ###
mr.stan.palleja <- stan(model, chains=4, iter=10000, data=palleja.data, warmup=1000)

# # Without asymptote
mr.stan.no.asym.palleja =  stan(model.no.asym, chains=4, iter=10000, data=palleja.data, warmup=1000)

bf.palleja <- compare_models(mr.stan.palleja, mr.stan.no.asym.palleja)


# 
# Produces plots of model fits
# Liam Shaw, 20th November 2017
library(easyGgplot2)
library(dplyr)
library(tidyr)
library(ggplot2)
library(rstan)
library(bridgesampling)
library(reshape2)
cat("libraries loaded")

# Set ggplot2 theme
theme_set(theme_bw() + theme(panel.margin = grid::unit(0, "lines")))

# Load previously fitted models

### PLOTTING FITS ####
makePlot <- function(data.df, stan.model, asymptote=TRUE, model.colour='grey', ylimits=c(-5, 8), title=""){
  plot(data.df$times, data.df$concs, pch=19, xlab="Months", ylab="Scaled diversity displacement", ylim=ylimits,
       cex.lab=1.3, cex.axis=1.3, main=title)
  times <- seq(0, 12, 0.1)
  stan.model.ex <- extract(stan.model)
  points <- matrix(nrow=1000, ncol=length(times))
  for(i in seq(1,1000)){
    if (asymptote==TRUE){
      points[i,] <- ssfolAsym(D=stan.model.ex$D[i], phi1=stan.model.ex$phi1[i], 
                              phi2=stan.model.ex$phi2[i], Asym=stan.model.ex$Asym[i], times)
    }
    else{
      points[i,] <- ssfol(D=stan.model.ex$D[i], phi1=stan.model.ex$phi1[i], 
                          phi2=stan.model.ex$phi2[i],times)
    }
  }
  # Add zero line
  abline(a=0,b=0, lwd=3, lty=2, col='grey')
  
  
  # Add 2.5%, median, 97.5% lines
  points.lower <-  apply(points, function(x) quantile(x, probs = 0.025), MARGIN=2)
  points.median <- apply(points, function(x) quantile(x, probs = 0.5), MARGIN=2)
  points.upper <- apply(points, function(x) quantile(x, probs=0.975), MARGIN=2)
  points(times, points.median, type='l', col=model.colour, lwd=2)
  points(times, points.lower, type='l', col=model.colour, lwd=2, lty=2)
  points(times, points.upper, type='l', col=model.colour, lwd=2, lty=2)
  
  # add data points on top
  points(data.df$times, data.df$concs, pch=19)
  
}

pdf('palleja-model-fit.pdf', width=14)
par(mfrow=c(1,2))
makePlot(palleja.data, mr.stan.no.asym.palleja, model.colour = 'red', asymptote = FALSE, ylimits = c(-2, 4), title="Model with return to initial equilibrium")
makePlot(palleja.data, mr.stan.palleja, model.colour = 'red', asymptote = TRUE, ylimits = c(-2, 4), title = "Model with change of equilibrium")
dev.off()

# Extract parameters
palleja.parameters <- extract(mr.stan.palleja)
palleja.no.asym.parameters <- extract(mr.stan.no.asym.palleja)

probs <- c(0.5, 0.025, 0.975)
quantile(palleja.parameters$D, probs = probs)
quantile(palleja.parameters$phi1, probs=probs)
quantile(palleja.parameters$phi2, probs=probs)
quantile(palleja.parameters$Asym, probs=probs)
```
