## Supplementary material for "Modelling microbiome recovery after antibiotics using a stability landscape framework": Supplementary-Text-1.pdf

### Main dataset: Zaura et al. (2015)

Zaura et al. (1) conducted a study on the long-term effect of antibiotics on the gut and oral microbiomes, where individuals were randomly assigned to one of five treatment groups: placebo, clindamycin, ciprofloxacin, minocycline, or amoxicillin (Table 1). Samples were collected at baseline, after treatment, one month, two months, four months, and one year. All treatments were administered for at most $\tau = 10$ days (150 mg clindamycin four times a day for ten days; 500 mg ciprofloxacin twice a day for ten days; 250 mg amoxicillin three times daily for seven days; 100mg minocycline twice daily for five days) and longitudinal faecal and saliva samples collected until $T = 1$ year afterwards i.e. $\frac{\tau}{T} \sim 0.027 \ll 1$. Therefore, the approximation of the antibiotics as an impulse perturbation (i.e. of negligible duration) should be valid. Samples underwent 16S rRNA gene amplicon sequencing, targeting the V5-V7 region (SRA Accession: SRP057504).

In order to reanalyze the data, we performed *de novo* clustering of sequences clustering into operational taxonomic units (OTUs) at 97% similarity with VSEARCH v1.1.1 (2) with chimeras removed against the 16S gold database (http://drive5.com/uchime/gold.fa). Taxonomy was assigned with RDP (3). For more details see Supplementary File 3, which contains more information and R code. The reanalyzed datasets are available as R phyloseq objects (Supplementary Files 4 and 5). We found no association between sequencing depth and timepoint.

We chose to use phylogenetic diversity as a metric of displacement (rather than e.g. richness or Shannon) because of its incorporation of phylogenetic information which should equate to a closer relationship with functional diversity of the microbiome. Phylogenetic diversity was calculated with the pd() function in the ‘picante’ R package v1.6-2 (4). Calculating this branch-weighted phylogenetic diversity requires a phylogeny, which we produced with FastTree v2.1.10 (5) after aligning 16S rRNA V5-V7 OTU sequences with Clustal Omega v1.2.1 (6). In order to compare uniform sequencing depths and obtain an estimate of error bars, we used bootstrapping by repeated subsampling of data to generate values for fitting the model. We used mean bootstrapped values ($n = 100$, sampling depth $r = 1\,000$) of phylogenetic diversity $d_i$ relative to the baseline phylogenetic diversity $d_0$ for each individual (Supplementary File 3), representing the displacement from equilibrium in our model ($\bar{d}_i = d_i - d_0$).

Bayesian model fitting was as described in the main text.

**A complex, individualized antibiotic response still allows a general model.** While it is not our intention to repeat a comprehensive description of the precise nature of the response for the different antibiotics (readers should refer to the original paper), we note some interesting qualitative observations from our reanalysis that highlight the complexity of the antibiotic response. We discuss here observations at the level of taxonomic family in the gut microbiomes of individuals taking ciprofloxacin or clindamycin (Supplementary Figure 1). While modelling these precise interactions is far beyond the scope of our model, our approach can still summarise the overall impact of this underlying complexity on the community as a whole.

Despite their different mechanisms of action, both clindamycin and ciprofloxacin caused a dramatic decrease in the Gram-negative anaerobes *Rikenellaceae*, which was most marked a month after the end of the course. However, for ciprofloxacin this decrease had already started immediately after treatment, whereas for clindamycin the abundance after treatment was unchanged in most participants. The different temporal nature of this response perhaps reflects the bacteriocidal nature of ciprofloxacin (7) compared to the bacteriostatic effect of clindamycin, although concentrations *in vivo* can produce bacteriocidal effects (8).

There were other clear differences in response between antibiotics. For example, clindamycin caused a decrease in the anaerobic Gram-positives *Ruminococcaceae* after a month, whereas ciprofloxacin had no effect. There was also an individualized response: ciprofloxacin led to dramatic increases in *Erysipelotrichaceae* for some participants, and for these individuals the increases coincided with marked decreases in *Bacteroidaceae*, suggesting the relevance of inter-family microbial interactions (Supplementary Figure 1).

Comparing relative abundances at the family level, there were few differences between community states of different treatment groups after a year. Equal phylogenetic diversity can be produced by different community composition, and this suggests against consistent trends in the long-term dysbiosis associated with each antibiotic. However, we did find that *Peptostreptococcaceae*, a member of the order *Clostridiales*, was significantly more abundant in the clindamycin group when compared to both the ciprofloxacin group and the placebo group separately ($p < 0.05$, Wilcoxon rank sum test). In a clinical setting, clindamycin is well-established to lead to an increased risk of a life-threatening infection caused by another member of *Clostridiales*: *Clostridium difficile* (9). Long-term reductions in diversity may similarly increase the risk of overgrowth of pathogenic species.

### Secondary dataset: Palleja et al. (2018)

Palleja et al. (10) gave twelve healthy individuals a four-day course of meropenem, gentamicin and vancomycin, and took samples at day 0, day 4, day 8, day 42, and day 180 which were sequenced with shotgun metagenomics. For fitting our model to this dataset, we downloaded rarefied mOTU relative abundances ('annotated.mOTU.rel_abund.rarefied.tsv') from the original author's website http://arumugamlab.sund.ku.dk/SuppData/Palleja_ et_al_2018_ABX. These represent inferred abundances of mOTUs based on the presence of ten single copy genes in the shotgun metagenomic data, rather than 16S rRNA gene based abundances. Rather than fully reanalyzing the dataset to calculate phylogenetic diversity, we wanted to see if this mOTU table could be used 'as is' to fit our model. We used the richness of each sample (number of species), scaling by an arbitrary factor of 50 ($x \to x/50$) in order to use the same parameter priors as for the Zaura et al. dataset which used phylogenetic diversity. See Supplementary File 6 for R code to reproduce results and Supplementary Figure 3.

Bayesian fitting was as in the main text (using Stan with the same models and parameter priors). We used 4 chains with a burn-in period of 1,000 iterations and 9,000 subsequent iterations, verifying that all chains converged ($\hat{R}$ = 1) and the effective sample size for each parameter was sufficiently large (*n_eff* > 1,000).
