## Supplementary material for "Modelling microbiome recovery after antibiotics using a stability landscape framework": Supplementary-Text-2.pdf

### 1 Supplementary text: Lotka-Volterra models

#### 1.1 Methods

The Lotka-Volterra equations are a simple, yet very powerful and popular, model of population dynamics in an ecosystem. They state that the rate of change of abundance of each species is determined by the abundance of that species and each other species, together with some parameters that are unique to each ecosystem. The Lotka-Volterra system is a subset of a generalised ecological model, in which interactions between species are restricted to pairwise interactions.[1] The mathematical form of the N-dimensional (i.e. for N interacting species) Lotka-Volterra equations can be seen in Equation 1.

$$\frac{dx_i}{dt} = \mu_i x_i(t) - x_i(t) \sum_{j=1}^{N} \alpha_{ij} x_j(t) \tag{1}$$

For $i = 1, 2, .., N$; where $x_i(t)$ is the abundance of species $i$ at time $t$, $\mu_i$ is the growth rate of species $i$ in the absence of any other effects, and the interaction term $\alpha_{ij}$ characterises the effect of species $j$ on species $i$, such that a positive value means one species activates another, a negative value means one species inhibits another and $\alpha_{ij} = 0$ indicates there is no interaction of species $j$ on species $i$ (note that $i$ could have an effect on $j$ unless $\alpha_{ji} = 0$ also).

To try and understand whether the same behaviour as the stability landscape model can be found using a Lotka-Volterra model, we can either try to understand this problem analytically or numerically. For reasons discussed further in Section 1.3, an analytic approach is currently unfeasible, and therefore we took a numerical approach to this problem. We modelled a three species ecosystem using the 3D Lotka-Volterra equations (equation 1 with $N = 3$). For a system like this, we added a perturbation term as done by Stein et al.[2] to give Equation 2.

$$\frac{dx_i}{dt} = \mu_i x_i(t) - x_i(t) \sum_{j=1}^{3} \alpha_{ij} x_j(t) + \varepsilon_i \left(\mathcal{H}(t-1.5) - \mathcal{H}(t-2.5)\right) x_i(t) \tag{2}$$

Where $\mathcal{H}(t)$ is the Heaviside step function, such that a perturbation of size $\varepsilon_i$ is applied to species $i$ from $t = 1.5$ to $t = 2.5$, modelling a day of antibiotics given 2 days after the start of the simulation. This timing was somewhat arbitrary, and the same behaviour in Section 1.2 was exhibited with a different choice of time for the perturbation.

Firstly, the analytical solution for all steady states was calculated (using Mathematica). This was done in the standard way by solving the set of simultaneous equations given by Equation 3.

$$\mu_i x_i - x_i \sum_{j=1}^{3} \alpha_{ij} x_j = 0 \tag{3}$$

This gives several potential steady states, the one with three non-zero values for $x_i, i = 1, 2, 3$ being

$$\frac{1}{-\alpha_{11}(\alpha_{22}\alpha_{33} + \alpha_{23}\alpha_{32}) + \alpha_{12}(\alpha_{21}\alpha_{33} - \alpha_{23}\alpha_{31}) - \alpha_{13}(\alpha_{21}\alpha_{32} + \alpha_{22}\alpha_{31})}(x_1^{\text{ss}}, x_2^{\text{ss}}, x_3^{\text{ss}}) \tag{4}$$

where

$$\begin{aligned}
x_1^{\text{SS}} &= \alpha_{12}\alpha_{23}\mu_3 - \alpha_{12}\alpha_{33}\mu_2 - \alpha_{13}\alpha_{22}\mu_3 + \alpha_{13}\alpha_{32}\mu_2 + \alpha_{22}\alpha_{33}\mu_1 - \alpha_{23}\alpha_{32}\mu_1 \\
x_2^{\text{SS}} &= -\alpha_{11}\alpha_{23}\mu_3 + \alpha_{11}\alpha_{33}\mu_2 + \alpha_{13}\alpha_{21}\mu_3 - \alpha_{13}\alpha_{31}\mu_2 - \alpha_{21}\alpha_{33}\mu_1 + \alpha_{23}\alpha_{31}\mu_1 \\
x_3^{\text{SS}} &= \alpha_{11}\alpha_{22}\mu_3 - \alpha_{11}\alpha_{32}\mu_2 - \alpha_{12}\alpha_{21}\mu_3 + \alpha_{12}\alpha_{31}\mu_2 + \alpha_{21}\alpha_{32}\mu_1 - \alpha_{22}\alpha_{31}\mu_1
\end{aligned}$$

Then we performed a grid search for parameters in which this created a feasible system with three-species coexistence ($x_i > 0 \ \forall i$) . That is to say that the growth rates were all positive ($\mu_i > 0 \ \forall i$) and the diagonal elements of the interaction matrix all negative ($\alpha_{ii} < 0 \ \forall i$), meaning that each species has a carrying capacity/maximum abundance, even in the absence of any other species. This is consistent with a biological system within a closed ecosystem, where unlimited abundance is not possible. The ranges used for each parameter were in line with models fitted to real biological data. [2]

For each of these parameter sets at the above fixed point, the Jacobian was calculated to determine whether the fixed point was stable. We are investigating the effect of perturbations on stable ecosystems, otherwise this behaviour could be easily found with unstable steady states with coexistence of three species. A steady state is stable if all eigenvalues of the Jacobian have a negative real part. [3] For all parameter sets with a stable steady state of three species coexisting, we then determined which parameter sets had another feasible steady state (with fewer than 3 species coexisting) and tested for that fixed points' stability if it does exist. If such parameter sets can be found, we should expect that a perturbation exists that changes the system from having 3 species coexisting to fewer than 3, i.e. causing a decrease in species diversity.

Once such a parameter set with these properties exists, a perturbation to move from the three species state to the lower diversity state could be calculated as being proportional to the difference between the position of the two states, as in Equation 5.

$$\varepsilon_i = K(x_i^{\text{altSS}} - x_i^{\text{SS}}) \qquad i = 1, 2, 3 \tag{5}$$

Where $K$ can be varied and the perturbation simulated in order to find the value that sufficiently moves the system to the alternative stable fixed point. If one wants to exclude the possibility of the antibiotic activating a certain species rather than inhibiting or having no effect, enforcing $\varepsilon_i < 0$ in the above equation still gives a suitable perturbation (though for a possibly different value of $K$). This provides just one possible perturbation that changes the system, others are certainly possible and can be found by a general grid search on all $\varepsilon_i$ for each set of model parameters.

#### 1.2 Results

For one set of growth parameters (in this case $\boldsymbol{\mu} = (0.9, 1, 1)$), searching over $5^9 = 1,953,125$ parameter sets (for the elements of the communits matrix $\alpha_i$), $93,155$ systems with a feasible 3 species coexistence were found, with just 74 sets possessing both stability in the three species coexistence and in one other fixed point (parameters for each of these are shown in Section 1.4). Figure 1 illustrates one of these sets, exhibiting the expected behaviour when perturbed. The direction and magnitude of potential perturbation vector (i.e. each species' susceptibility to the antibiotics) required to make such behaviour varied for each system, with one example discussed previously.

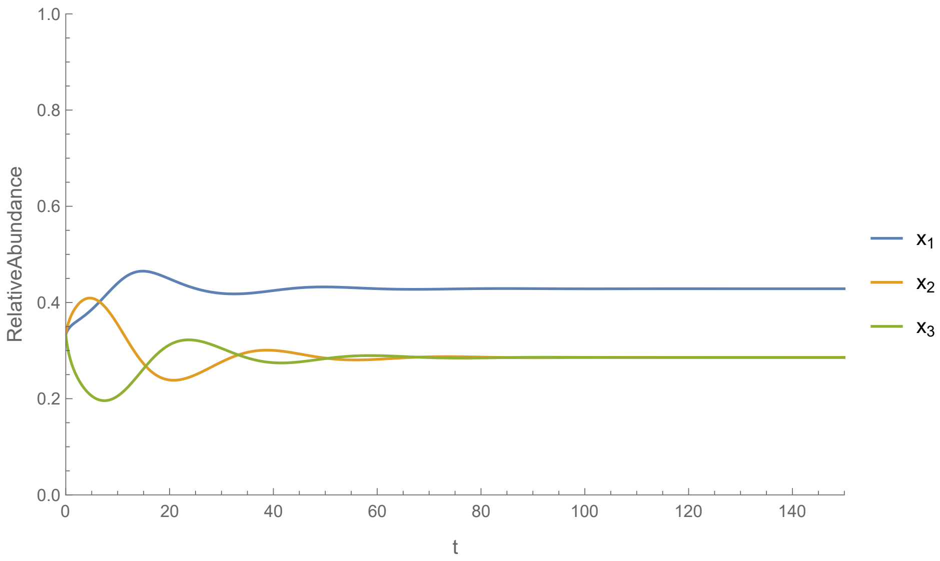


(a) No perturbation

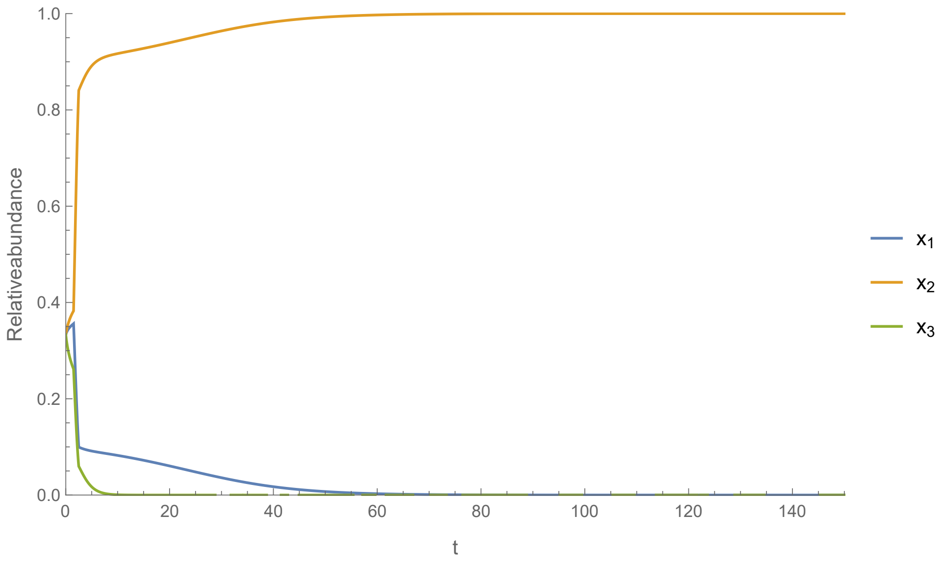


(b) Large perturbation

Figure 1: Dynamics of one Lotka-Volterra system. (a) shows that the coexistence of three-species is reached in absence of perturbations. (b) When perturbed, only one species survives, with $x_1$ and $x_3$ going to zero.

The gut microbiome is stable when subject to small perturbations[4], which are encountered constantly in human biology, but can have its makeup altered when subject to particularly large perturbations. Similarly, in Figure 2, a smaller perturbation (half the size of that in Figure 1) causes a change in the abundances of each species in the above system but over time, the system returns to its original state. Therefore, the exact antibiotic and how it interacts with the existing microbiota will effect whether the antibiotic treatment will 'permanently' change the makeup of the gut microbiome.

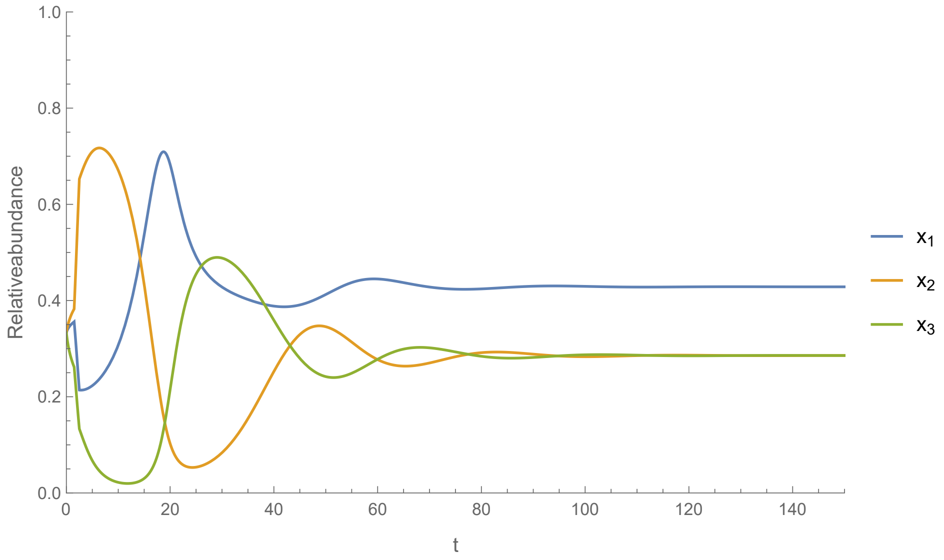


Figure 2: Small perturbation applied to Lotka-Volterra system from Figure 1, which returns the system to its original state of three coexisting species. This corresponds to an antibiotic or other perturbation that inhibits species 1 and 3 half a much as in Figure 1

It should also be noted that the 'step' perturbation in Equation 2 can be substituted for an 'impulse' perturbation, as in the stability landscape model, without a change in the results (other than the necessary value for parameter $K$ in Equation 5). This suggests the exact manifestation of the perturbation is perhaps not incredibly relevant, at least for the Lotka-Volterra model. One exception might be when looking at long term use of antibiotics. Theoretically even smaller perturbations could then result in the microbiome never being in equilibrium, with perturbations as in 1 occurring before the effect of the last one have been forgotten by the system. This could also lead to an accumulation in perturbations that moves the system to the alternative state, but further modelling would be required to see whether this is the case.

#### 1.3 Discussion

We showed that it is possible in a three species Lotka-Volterra model to exhibit the same behaviour as that of the two-state stability landscape model. We searched $5^9 = 1,953,125$ randomly chosen parameter sets and identified $93,155$ systems with a stable state (fixed point) where 3 species coexisted. Of these, 74 had an additional fixed point with fewer than 3 species, which the system could reach if perturbed sufficiently. The observation that 0.079% of three-species

Lotka-Volterra systems exhibit the behaviour required by the two-state model suggests that the stability landscape model is unrealistic for small numbers of species, which is as expected because the stability landscape mode assumes diversity (the state of the microbiome) is a continuous variable rather than a clearly discrete one with only certain permitted values. However, in principle this behaviour is possible for some parameter sets even at low $n$.

For larger numbers of species, recent work in theoretical ecology gives a strong justification for the generalisability of this behaviour to higher dimensions. It has recently been shown that as the number of species $n$ increases, the number of fixed points which are stable increases independently of population size [5], and the proportion of simulations from random parameters that have multiple fixed points also increases: with $n = 400$, this proportion is $> 97\%$ [35]. This suggests that the overwhelming majority of mathematically possible systems at relevant numbers of species exhibit multible fixed points; we suggest the likelihood is that the fraction of biologically possible systems exhibiting this behaviour is even higher. Furthermore, the Lotka-Volterra model undergoes a phase transition from a system with a unique fixed point (UFP) to one with multiple attractors (MA); when resource competition is incorporated into the model  a more realistic assumption in the case of the human microbiome  all fixed points become either stable or marginally stable Bunin [6]. The gut microbiome is an ecosystem of hundreds of species in the presence of resource competition. From this assumption, we can therefore infer from this theoretical work that: it exists beyond the UFP phase; its community composition will be history-dependent; and perturbations will lead to transitions between the multiple possible stable states. This high level behaviour is then captured by the two-state stability-landscape model using only the simple measure of diversity.

Further to the long term decrease in diversity compared to the initial state, it can be seen in the Figure 1 that the perturbation has a partially delayed effect on species diversity. Depending on the size of the perturbation, the lowest species diversity may not be seen until a significant time after the perturbation occurs. This is inline with the behaviour of the stability landscape model by Shaw et al. [7] and the observation made by Zaura et al. [8], where their dataset showed that the lowest diversity was observed one month after the antibiotic perturbation, rather than immediately after.

#### 1.4 Table of parameter values

The parameters were taken as follows: $\boldsymbol{\mu} = (0.9, 1, 1)$, $\alpha_{ij} \in \{-1, -0.5, 0, 0.5, 1\}$ for $i \neq j$, $\alpha_{ii} \in \{-2, -1.5, -1, -0.5, 0\}$. Table 1 shows the 74 sets of parameters which gave the desired behaviour from those simulated.

| $\alpha_3$ | $\alpha_4$ | $\alpha_5$ | $\alpha_6$ | $\alpha_7$ | $\alpha_8$ | $\alpha_9$ |
|---|---|---|---|---|---|---|
| -1. | -1. | -0.5 | -0.5 | 0. | -1. | -1.5 |
| -1. | -1. | -0.5 | -0.5 | 0.5 | -1. | -2. |
| -1. | -1. | -0.5 | -0.5 | 0.5 | -1. | -1.5 |
| -1. | -1. | -0.5 | -0.5 | 1. | -1. | -2. |
| -1. | -1. | -0.5 | -0.5 | 1. | -1. | -1.5 |
| -1. | -1. | -0.5 | 0. | 0. | -1. | -2. |
| -0.5 | -1. | -0.5 | 0. | 0. | -1. | -1. |
| -1. | -1. | -0.5 | 0. | 0. | -1. | -1.5 |
| -1. | -1. | -0.5 | 0. | 0.5 | -1. | -2. |
| -0.5 | -1. | -0.5 | 0. | 0.5 | -1. | -1. |
| -1. | -1. | -0.5 | 0. | 0.5 | -1. | -1.5 |
| -1. | -1. | -0.5 | 0. | 1. | -1. | -2. |
| -0.5 | -1. | -0.5 | 0. | 1. | -1. | -1. |
| -1. | -1. | -0.5 | 0.5 | 0. | -1. | -2. |
| -1. | -1. | -0.5 | 0.5 | 0. | -1. | -1.5 |
| -1. | -1. | -0.5 | 0.5 | 0.5 | -1. | -2. |
| -1. | -1. | -0.5 | 0.5 | 1. | -1. | -2. |
| -1. | -1. | -0.5 | 1. | 0. | -1. | -2. |
| -0.5 | -1. | -0.5 | 0.5 | 0. | -1. | -1. |
| -1. | -1. | -0.5 | 1. | 0. | -1. | -1.5 |
| -1. | -1. | -0.5 | 1. | 0.5 | -1. | -2. |
| -0.5 | -1. | -0.5 | 0.5 | 0.5 | -1. | -1. |
| -1. | -1. | -0.5 | 0. | 0. | -1. | -2. |
| -0.5 | -1. | -0.5 | 0. | 0. | -1. | -1. |
| -0.5 | -1. | -0.5 | 0. | 0.5 | -1. | -1.5 |
| -1. | -1. | -0.5 | 0. | 0.5 | -1. | -2. |
| -0.5 | -1. | -0.5 | 0. | 0.5 | -1. | -1. |
| -0.5 | -1. | -0.5 | 0. | 1. | -1. | -2. |
| -0.5 | -1. | -0.5 | 0. | 1. | -1. | -1.5 |
| -1. | -1. | -0.5 | 0. | 1. | -1. | -2. |
| -0.5 | -1. | -0.5 | 0. | 1. | -1. | -1. |
| -0.5 | -1. | -0.5 | 0.5 | 0. | -1. | -1.5 |
| -1. | -1. | -0.5 | 1. | 0. | -1. | -2. |
| -0.5 | -1. | -0.5 | 0.5 | 0. | -1. | -1. |
| -0.5 | -1. | -0.5 | 0.5 | 0.5 | -1. | -2. |
| -0.5 | -1. | -0.5 | 0.5 | 0.5 | -1. | -1.5 |
| -1. | -1. | -0.5 | 1. | 0.5 | -1. | -2. |

| $\alpha_3$ | $\alpha_4$ | $\alpha_5$ | $\alpha_6$ | $\alpha_7$ | $\alpha_8$ | $\alpha_9$ |
|---|---|---|---|---|---|---|
| -0.5 | -1. | -0.5 | 0.5 | 0.5 | -1. | -1. |
| -0.5 | -1. | -0.5 | 0.5 | 1. | -1. | -2. |
| -0.5 | -1. | -0.5 | 0.5 | 1. | -1. | -1.5 |
| -0.5 | -1. | -0.5 | 1. | 0. | -1. | -2. |
| -0.5 | -1. | -0.5 | 1. | 0. | -1. | -1.5 |
| -0.5 | -1. | -0.5 | 1. | 0. | -1. | -1. |
| -0.5 | -1. | -0.5 | 1. | 0.5 | -1. | -2. |
| -0.5 | -1. | -0.5 | 1. | 0.5 | -1. | -1.5 |
| -0.5 | -1. | -0.5 | 1. | 0.5 | -1. | -1. |
| -0.5 | -1. | -0.5 | 1. | 1. | -1. | -2. |
| -0.5 | -1. | -0.5 | 1. | 1. | -1. | -1.5 |
| 0. | -1. | -0.5 | 0.5 | 0. | -1. | -0.5 |
| 0. | -1. | -0.5 | 1. | 0. | -1. | -1. |
| 0. | -1. | -0.5 | 0.5 | 0.5 | -1. | -1. |
| 0. | -1. | -0.5 | 1. | 0.5 | -1. | -2. |
| 0. | -1. | -0.5 | 0.5 | 0.5 | -1. | -0.5 |
| 0. | -1. | -0.5 | 1. | 0.5 | -1. | -1. |
| 0. | -1. | -0.5 | 0.5 | 1. | -1. | -1.5 |
| 0. | -1. | -0.5 | 0.5 | 1. | -1. | -1. |
| 0. | -1. | -0.5 | 1. | 1. | -1. | -2. |
| 0. | -1. | -0.5 | 0.5 | 1. | -1. | -0.5 |
| 0. | -1. | -0.5 | 1. | 1. | -1. | -1. |
| 0. | -1. | -0.5 | 0.5 | 0. | -1. | -0.5 |
| 0. | -1. | -0.5 | 1. | 0. | -1. | -1. |
| 0. | -1. | -0.5 | 1. | 0. | -1. | -0.5 |
| 0. | -1. | -0.5 | 0.5 | 0.5 | -1. | -1. |
| 0. | -1. | -0.5 | 1. | 0.5 | -1. | -2. |
| 0. | -1. | -0.5 | 1. | 0.5 | -1. | -1.5 |
| 0. | -1. | -0.5 | 0.5 | 0.5 | -1. | -0.5 |
| 0. | -1. | -0.5 | 1. | 0.5 | -1. | -1. |
| 0. | -1. | -0.5 | 1. | 0.5 | -1. | -0.5 |
| 0. | -1. | -0.5 | 0.5 | 1. | -1. | -1. |
| 0. | -1. | -0.5 | 1. | 1. | -1. | -2. |
| 0. | -1. | -0.5 | 1. | 1. | -1. | -1.5 |
| 0. | -1. | -0.5 | 0.5 | 1. | -1. | -0.5 |
| 0. | -1. | -0.5 | 1. | 1. | -1. | -1. |
| 0.5 | -1. | -0.5 | 1. | 1. | -1. | -0.5 |

Table 1: Table of parameter values for the community matrix $\alpha_i$ which exhibited multiple stable fixed points, with $\alpha_i = -0.5 \quad i = 1, 2$ in all cases excluded for ease of presentation. All had one point with three species coexisting, and another with just one species surviving. $\boldsymbol{\mu} = (0.9, 1, 1)$ for these simulations. $\alpha_{ij} \in \{-1, -0.5, 0, 0.5, 1\} \quad \text{for} \quad i \neq j, \qquad \alpha_{ii} \in \{-2, -1.5, -1, -0.5, 0\}$

[3] Marc R Roussel. Stability analysis for odes.

[4] Vanni Bucci, Serena Bradde, Giulio Biroli, and Joao B Xavier. Social interaction, noise and antibiotic-mediated switches in the intestinal microbiota. *PLoS computational biology*, 8(4):e1002497, 2012.

[5] Theo Gibbs, Jacopo Grilli, Tim Rogers, and Stefano Allesina. The effect of population abundances on the stability of large random ecosystems. *arXiv preprint arXiv:1708.08837*, 2017.

[6] Guy Bunin. Ecological communities with lotka-volterra dynamics. *Physical Review E*, 95(4):042414, 2017.

[7] Liam P Shaw, Chris P Barnes, A Sarah Walker, Nigel Klein, and Francois Balloux. A perturbation model of the gut microbiome's response to antibiotics. *bioRxiv*, page 222398, 2017.

[8] Egija Zaura, Bernd W Brandt, M Joost Teixeira de Mattos, Mark J Buijs, Martien PM Caspers, Mamun-Ur Rashid, Andrej Weintraub, Carl Erik Nord, Ann Savell, Yanmin Hu, et al. Same exposure but two radically different responses to antibiotics: resilience of the salivary microbiome versus long-term microbial shifts in feces. *MBio*, 6(6):e01693–15, 2015.
