## Supplementary figures and images for "Modelling microbiome recovery after antibiotics using a stability landscape framework"

### Supplementary-Figure-1.pdf

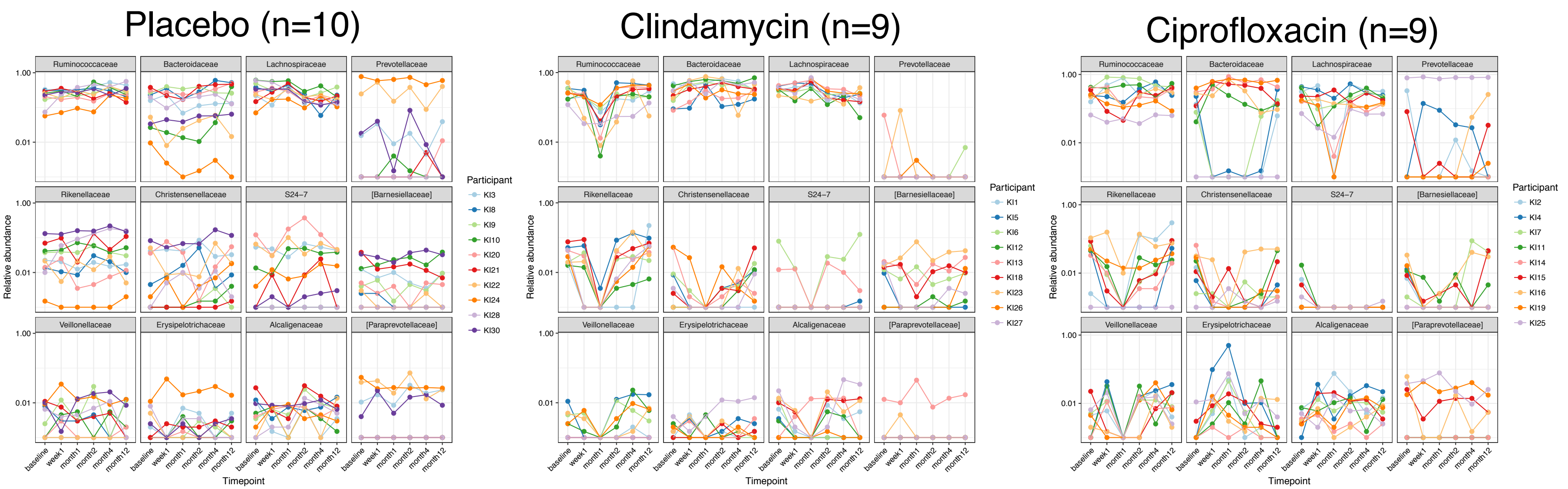

### Supplementary-Figure-2.pdf

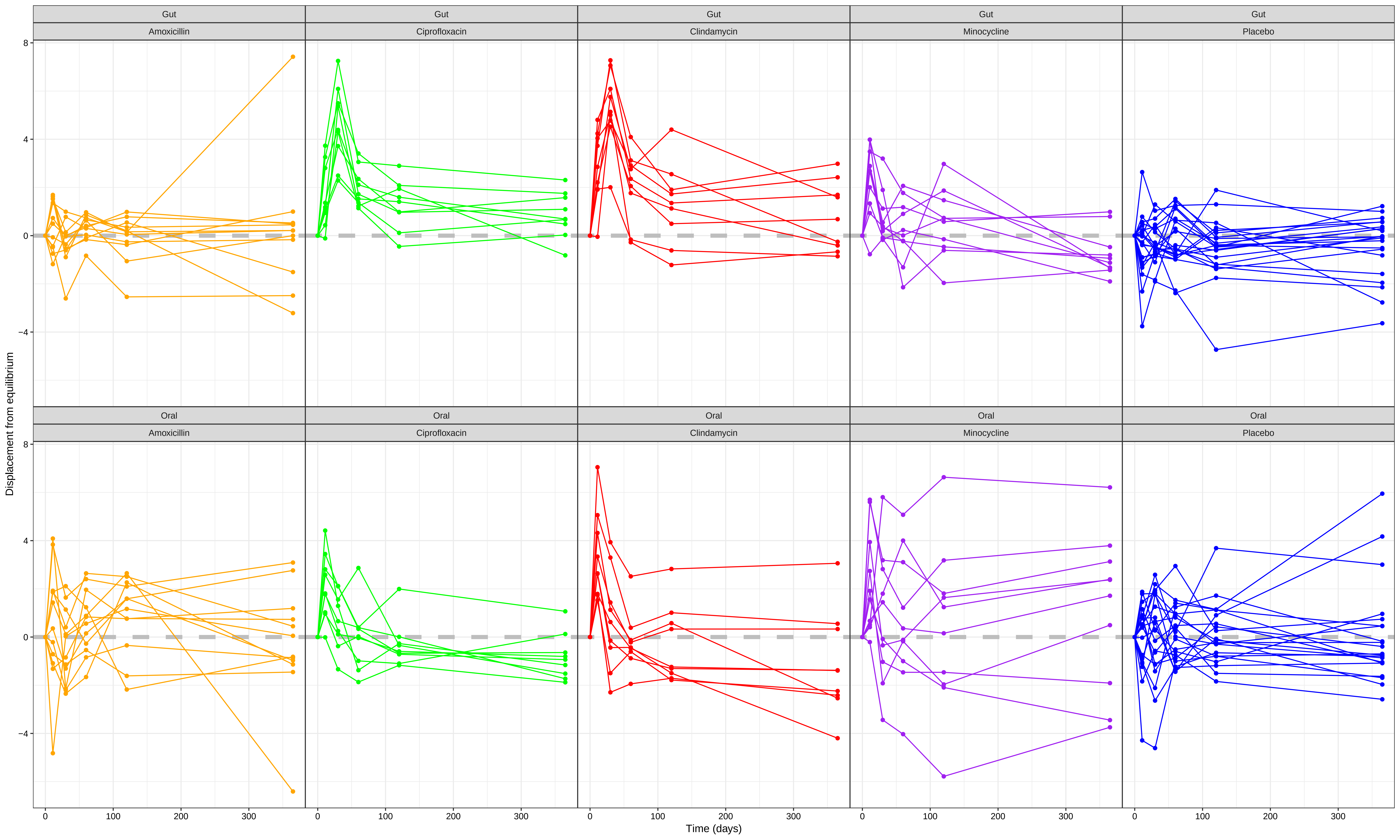

### Supplementary-Figure-3.pdf

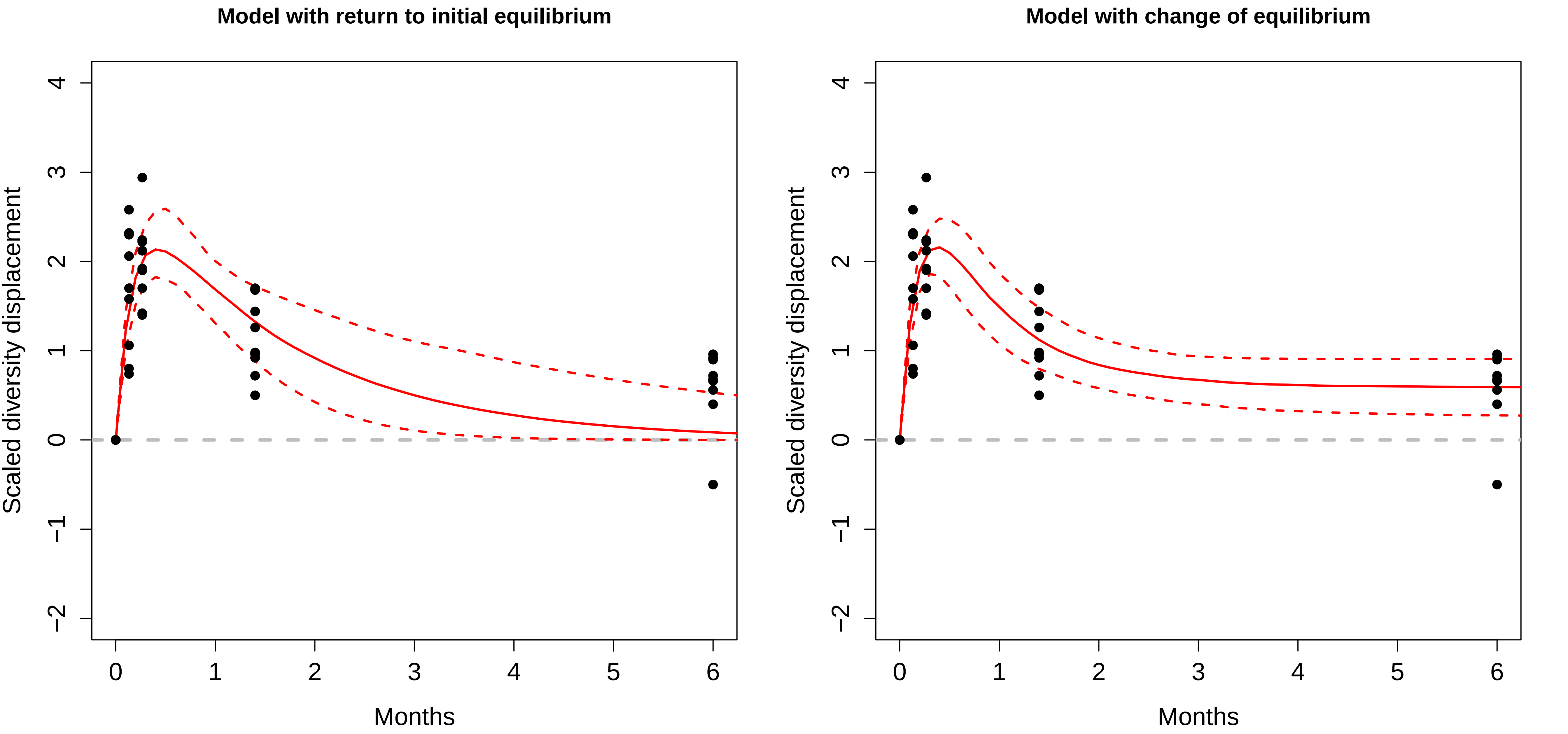
